# A major endogenous glycosidase mediating quercetin uptake in *Bombyx mori*

**DOI:** 10.1101/2023.08.02.551622

**Authors:** Ryusei Waizumi, Chikara Hirayama, Shuichiro Tomita, Tetsuya Iizuka, Seigou Kuwazaki, Akiya Jouraku, Takuya Tsubota, Kakeru Yokoi, Kimiko Yamamoto, Hideki Sezutsu

**Author notes:** Correspondence: Ryusei Waizumi.

## Abstract

Quercetin is a common plant flavonoid that is involved in herbivore–plant interactions. Mulberry silkworms (domestic silkworm, *Bombyx mori*, and wild silkworm, *Bombyx mandarina*) uptake quercetin from mulberry leaves and accumulate the metabolites in the cocoon, thereby improving its protective properties. Here we identified and characterized a glycosidase, named LPH-like quercetin glycoside hydrolase 1 (LQGH1), that initiates quercetin metabolism in the domestic silkworm. LQGH1 is expressed in the midgut where it mediates quercetin uptake by deglycosylating the three most common quercetin glycosides present in mulberry leaf: rutin, quercetin-3-*O*-malonyl-glucoside, quercetin-3-*O*-glucoside. Despite being located in an unequal crossing-over hotspot, *LQGH1* is conserved in some species in clade Macroheterocera, including the wild silkworm, indicating the adaptive significance of quercetin uptake. *LQGH1* is important also in breeding: defective mutations of *LQGH1*, which result in discoloration of the cocoon and increased silk yield, are homozygously conserved in 27 of the 32 Japanese white-cocoon domestic silkworm strains and 12 of the 30 Chinese ones we investigated.

## Introduction

Quercetin (3,3’,4’,5,7-pentahydroxyflavone) is a flavonoid abundantly found in a wide variety of plants[1]. Previous studies have found oviposition and feeding stimulant activity of quercetin glycosides on lepidopteran, orthopteran and coleopteran insects [2,3,4,5,6]. These suggest that quercetin is widely ingested by insects. Quercetin ingestion by insects is likely not merely a consequence of the identification and feeding on host plants; it may have adaptive significance. In the common blue butterfly (*Polyommatus icarus*), which sequesters quercetin-3-*O*-galactoside in its wings, flavonoid content is high in the female insects than in the males and positively correlated with female sex attraction [7,8,9]. The yellow pigment in the wings of a grasshopper (*Dissosteira carolina*), which possibly contributes to their camouflage in plants, is the result of sequestration of quercetin-3-*O*-glucoside (isoquercitrin, Q3G)[10]. Although these studies strongly emphasize the significance of quercetin in herbivore–plant interactions, the molecular mechanisms of quercetin metabolism in insects are poorly understood.

Mulberry silkworms (domestic silkworm, *Bombyx mori*, and wild silkworm, *Bombyx mandarina*) accumulate various compounds, including quercetin glucosides, kaempferol glucosides, and carotenoids, in their silk glands and colored cocoons (S1 Fig) [11,12,13]. Quercetin glucosides stored in the body and in the cocoon are reported to have antioxidant, ultraviolet-protective, and antibacterial properties [14,15,16]. Quercetin is found in the leaves of the mulberry tree (*Morus alba*), the sole food source of the mulberry silkworm, as a series of glycosides formed by glycosylation at the 3-*O* position. The three most common quercetin glycosides in mulberry are quercetin-3-*O*-rutinoside (rutin), quercetin-3-*O*-malonyl-glucoside (Q3MG) and Q3G, which account for 71%–80% of the total flavonol content in the leaves[17].

The color of the cocoon of the domestic silkworm has been diversified through breeding, which indicates that the kinds and amounts of flavonoids that accumulate in the cocoons differ between strains [11,12]. Particularly, cocoons containing high flavonoid concentrations express a yellow-green color and are known as “green cocoons”. Because accumulation in the cocoon is the end step of flavonoid metabolism, forward-genetic analysis focused on cocoon flavonoid content can reveal the genes involved in each step of flavonoid metabolism. Indeed, several loci associated with flavonoid metabolism have already been identified through this approach: the *Green b* locus, which encodes a uridine 5’-diphospho-glucosyltransferase with a rare enzymic activity glycosylating the 5-*O* position of quercetin[14], and the *New Green Cocoon* (*Gn*) locus, which encodes clustered sugar transporters presumed to import quercetin glucosides from the hemolymph to the silk gland[18]. However, these findings explain only a part of the process from quercetin uptake to the final accumulation of its metabolites in the cocoon. Elucidating the steps of flavonoid metabolism in the silkworm can provide valuable insights into the understanding of herbivore–plant interactions.

Here, we performed a quantitative trait locus (QTL) analysis focused on cocoon flavonoid content in the domestic silkworm. We identified a novel locus, *Green d* (*Gd*), that is associated with cocoon flavonoid content. Within the locus, we identified a glycosidase gene, *LPH-like quercetin glycoside hydrolase 1* (*LQGH1*), that mediates quercetin uptake into the midgut cells by deglycosylating mulberry leaf-derived quercetin glycosides. Genetic dissections of the novel gene revealed the contribution of the gene to improvement of the cocoon in the commercial context through breeding.

## Results

### QTL analysis identified a novel locus associated with flavonoid content in cocoon

To identify genes involved in quercetin metabolism by means of a forward-genetics approach, we prepared a green-cocoon strain (p50; alias: Daizo) and a white-cocoon strain (J01; alias: Nichi01). The two strains exhibited a distinct difference in cocoon color and flavonoid content (Fig 1A and 1B). A gradated range of cocoon colors in their F2 intercrossing offspring implied that the genetic differences between the two strains associated with flavonoid content were composite (Fig 1C). Therefore, we conducted a QTL analysis, which allows for simultaneous identification of multiple genetic loci involved in a phenotype of interest. The flavonoid content of the cocoons of the F2 population were scored by an absorbance-based method according to a previous report[19]. The QTL analysis was performed by using the phenotypic data of 102 individuals and 1038 genetic markers obtained from double-digest restriction-associated DNA sequencing data[20] (S1 Table). From the composite interval mapping, we identified three significant QTLs for cocoon flavonoid content on chromosomes 15, 20, and 27 (Fig 1C). Previous linkage studies have suggested a locus, named *Green c* (*Gc*) and associated with the yellow-green color of cocoons, at an unknown position on chromosome 15 [21,22]. The QTL on chromosome 27 was presumed to correspond to *Gn*[18]. Since no green cocoon-associated locus on chromosome 20 has yet been reported, we named that locus *Green d* (*Gd*). The contribution of the *Gd* locus to cocoon flavonoid content was the second largest of the three QTLs, with a percentage of phenotypic variation explained by each QTL (PVE) value of 24.57% (Fig 1C, S2 Table). The 95% Bayes credible interval of the *Gd* locus was 7,980,189–10,504,065 bp. The nearest marker to *Gd* was located at 10,265,033 bp, with a logarithm of odds (LOD) score of 19.99. The PVE values of the QTLs on chromosomes 15 and 27 were 7.04% and 56.05%, respectively. Significant additive effects were detected between the QTLs on chromosomes 15 and 27, and between those on chromosomes 20 and 27, and the PVEs were 1.93% and 5.47%, respectively (S2 Table).

**Fig 1.**
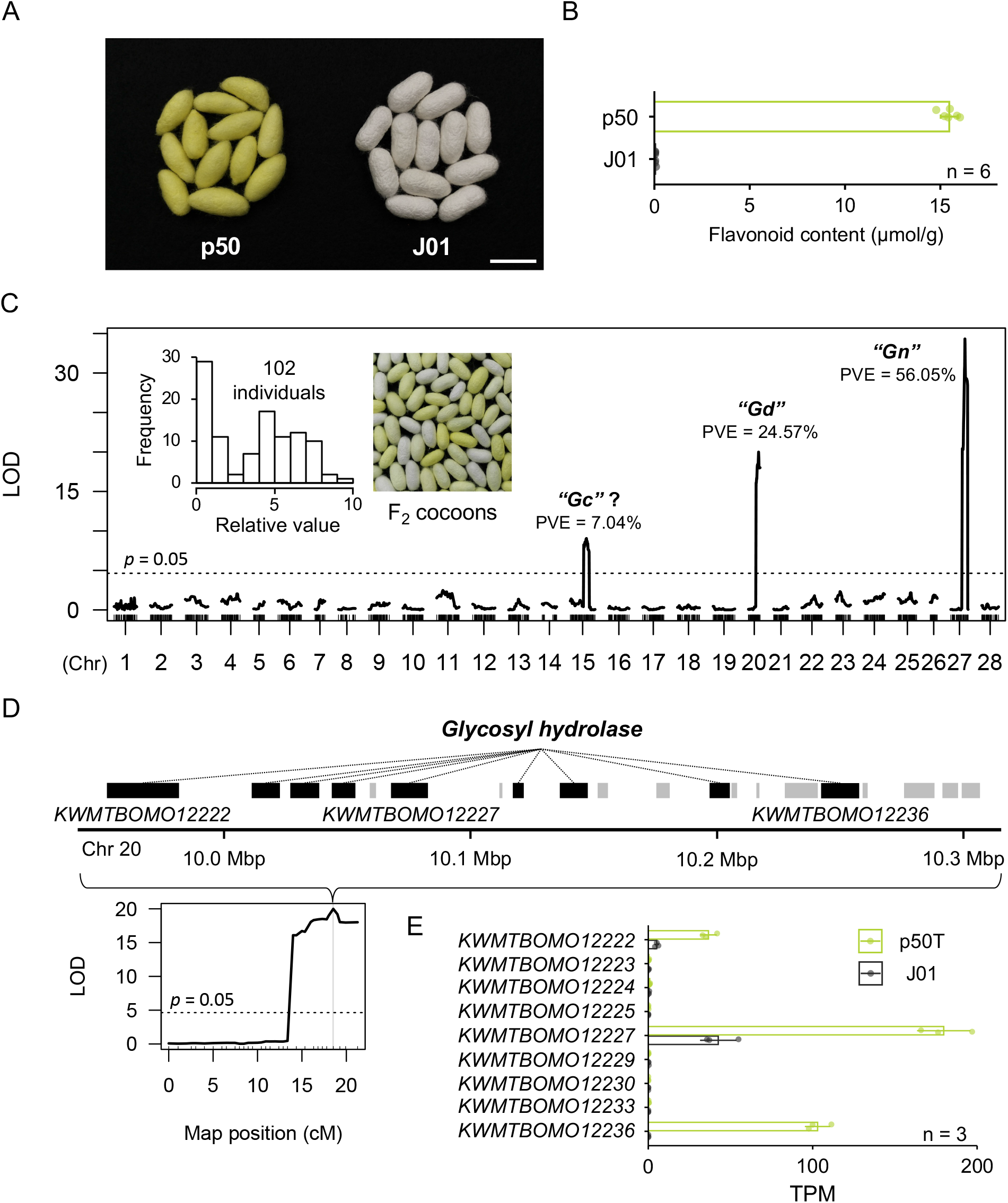
Quantitive trait loci (QTLs) associated with cocoon flavonoid content. (A) Photographs of representative cocoons of the p50 and J01 silkworm strains. Bar = 30 mm. (B) Flavonoid content of the cocoon of p50 and J01. Data are means ± SD. (C) QTL analysis for cocoon flavonoid content. The horizontal dotted line indicates the threshold of the permutation test (trials = 1000). Phenotype scores on the frequency distribution are relative to a maximum measurement of 10. LOD, logarithm of odds; PVE, percentage of phenotypic variation explained by each QTL. According to a previous study[21], the *Gc* locus is located on chromosome 15, but the detailed genetic or physical position is unknown. Therefore, it cannot be concluded that the QTL peak on chromosome 15 identified here corresponds to the *Gc* locus. (D) Modelled genes present within the *Gd* locus of the p50T genome assembly. The genes annotated as encoding a glycosyl hydrolase are highlighted in black; their IDs are *KWMTBOMO12222*, *KWMTBOMO12223*, *KWMTBOMO12224*, *KWMTBOMO12225*, *KWMTBOMO12227*, *KWMTBOMO12229*, *KWMTBOMO12230*, *KWMTBOMO12233*, and *KWMTBOMO12236* from the upstream side. (E) Expression of the candidate *Gd* genes in the midgut of third-day final instar male larvae. Data are means ± SD. TPM, transcripts per million.

### Gd locus contains a glycosidase cluster

Using the genome assembly and gene models of p50T[23], we found on the *Gd* locus a cluster of nine genes encoding glycoside hydrolases (*KWMTBOMO12222–25*, *KWMTBOMO12227*, *KWMTBOMO12229*, *KWMTBOMO12230*, *KWMTBOMO12233*, and *KWMTBOMO12236*) (Fig 1D). In mammals, deglycosylation of quercetin glucosides by lactase/phlorizin hydrolase (LPH), a member of beta-glucosidase family 1, is a critical step in quercetin metabolism. LPH is expressed in intestinal epithelial cells and is anchored on the brush border membrane where it hydrolyzes flavonoid glycosides [24,25,26,27]. The resulting free quercetin aglycon is then passively absorbed into the intestinal cells due to its increased lipophilicity. We hypothesized that the glycosidases clustered within the *Gd* locus are involved in quercetin metabolism in the domestic silkworm, playing a role similar to that of LPH in mammals. To identify which of the *Gd* glycosidases act in the midgut, RNA-seq-based expression analysis was performed. Three of the candidate genes, *KWMTBOMO12222*, *KWMTBOMO12227*, and *KWMTBOMO12236*, were found to be strongly expressed in the midgut of p50T final instar larvae; the expression levels of these genes were significantly lower in the midgut of strain J01 (Fig 1E). Further examination of the expression profiles of the three genes with high expression in the midgut revealed that *KWMTBOMO12227* and *KWMTBOMO12236* were expressed specifically in the midgut, whereas *KWMTBOMO12222* was expressed most in the malpighian tubule (S2 Fig). Thus, we identified *KWMTBOMO12222*, *KWMTBOMO12227*, and *KWMTBOMO12236* as candidate *Gd* genes which are involved in quercetin metabolism in the domestic silkworm.

### Functional analysis of the candidate *Gd* genes by CRISPR-Cas9

To investigate the involvement of the candidate genes in the quercetin metabolism, we attempted to use a microinjection-mediated CRISPR-Cas9 system to establish p50T lineages in which the candidate genes had been knocked out. Although, we failed to establish knockout lineages for *KWMTBOMO12222* and *KWMTBOMO12236* due to the lethality of homozygous frame-shift mutations, we did manage to obtain two knockout lineages of *KWMTBOMO12227* with different types of frameshift mutations in exon 5. We designated the one with a 5-bp deletion as Δ*Gd1* and the other with a 2-bp deletion as Δ*Gd2* (Fig 2A). These mutations resulted in a premature stop codon at exon 5 and a shortened amino acid sequence length of KWMTBOMO12227 from 492 to 215 in Δ*Gd1* and to 216 in Δ*Gd2*. Further characterization revealed that compared to p50T the mutants produced discolored cocoons (Fig 2B). In addition, a reduction of fluorescence under ultraviolet irradiation, which is characteristic of the accumulation of the two major quercetin metabolites in silkworm (i.e., quercetin-5-*O*-glucoside and quercetin-5,4′-di-*O*-glucoside) [14,15,28], was observed in the midgut, hemolymph, and silk glands of the mutants (Fig 2C–2E), indicating reductions of flavonoid content in the mutant tissues. Although knockout of *KWMTBOMO12227* reduced the total flavonoid content in the cocoon by less than half that in the original p50T strain, it was still much larger than the effect predicted for the *Gd* locus in the QTL analysis (Fig 2F and S2 Table). Similar reductions were found in the midgut, hemolymph, and middle and posterior silk gland. Because the flavonoid content in the cocoon differed largely between the insects reared with an artificial diet and those reared with fresh mulberry leaves (Fig 1B and Fig 2F), the flavonoid content reductions in the mutants were confirmed in an experiment using fresh mulberry leaves (S3 Fig). Together, these results indicate that knockout of *KWMTBOMO12227* results in malfunction of quercetin uptake into the midgut.

**Fig 2.**
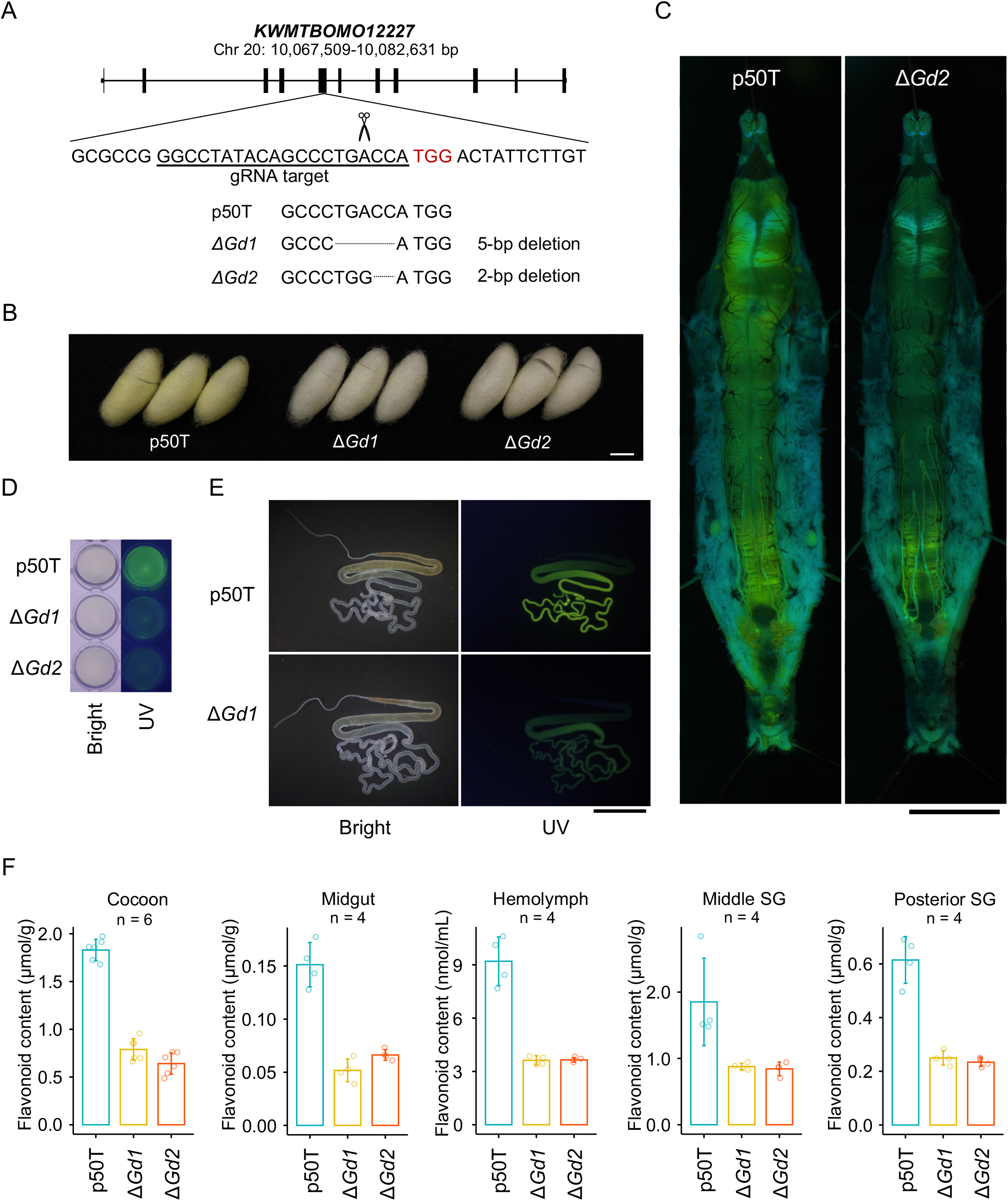
Reduction of flavonoid content in *KWMTBOMO12227-*knockout mutants. (A) Disrupted target sequences in *KWMTBOMO12227*. (B)–(E) Cocoons (B), midguts (C), hemolymph (D), and silk glands (E) of the p50T strain and the *KWMTBOMO12227* mutants. Organs and tissues were taken from sixth-day final instar female larvae. Bars = 10 mm. (F) Total flavonoid content of cocoons, organs, and tissues of the p50T strain and the *KWMTBOMO12227* mutant lineages. Data are means ± SD. All insects were reared on a commercial artificial diet. SG, silk gland.

### Characterization of the protein sequence of the glycosidase encoded by *KWMTBOMO12227*

The CRISPR-Cas9-mediated knockout analysis strongly suggested that *KWMTBOMO12227* encodes a glycosidase which plays a crucial role in quercetin uptake in the domestic silkworm. *KWMTBOMO12227* consists of 11 exons, encoding a total of 492 amino acids. The accuracy of the modeled sequence was confirmed using the RNA-seq data obtained from final instar larvae of strain p50T (S1 Text). KWMTBOMO12227 was found to have an amino acid sequence similar to that of LPH, but phylogenetic inference suggested that KWMTBOMO12227 is not orthologous to LPH; it has evolved through gene duplication after the divergence of Pyraloidea and Macroheterocera (including Noctuoidea, Lasiocampoidea, and Bombycoidea) (S1 Text). Mature LPH harbors a transmembrane domain in the C-terminus region with which it is anchored to the brush border membrane[25]. A hydrophobic transmembrane domain was also predicted in the first N-terminal 20 amino acid residues of KWMTBOMO12227 (S4 Fig), suggesting that the protein functions as a membrane-anchored glycosidase.

### LQGH1 mediates quercetin uptake by hydrolysis of quercetin glycosides

Together, our knockout analysis and sequence characterization suggested that the glycosidase encoded by *KWMTBOMO12227* is anchored on the apical surface of epithelial cells and mediates the uptake of mulberry-derived quercetin by deglycosylation of quercetinglycosides in the midgut of the silkworm. However, some accumulation of flavonoids in the midgut was still observed in the knockout mutants (Fig 2F). This may be due to the diversity of quercetin glycosides in mulberry leaves and the substrate specificity of KWMTBOMO12227. To confirm whether *KWMTBOMO12227* is involved in deglycosylation of quercetin glycosides, we investigated the effect of knocking out the gene on the hydrolytic activity of midgut tissue on the three major quercetin glycosides in mulberry leaf: rutin, Q3MG, and Q3G (Fig 3A). To also examine the localization of KWMTBOMO12227, the midgut homogenate and the insoluble material fraction of that, including cell membranes, isolated from the homogenate by centrifugation, were used separately for the measurements.

**Fig 3.**
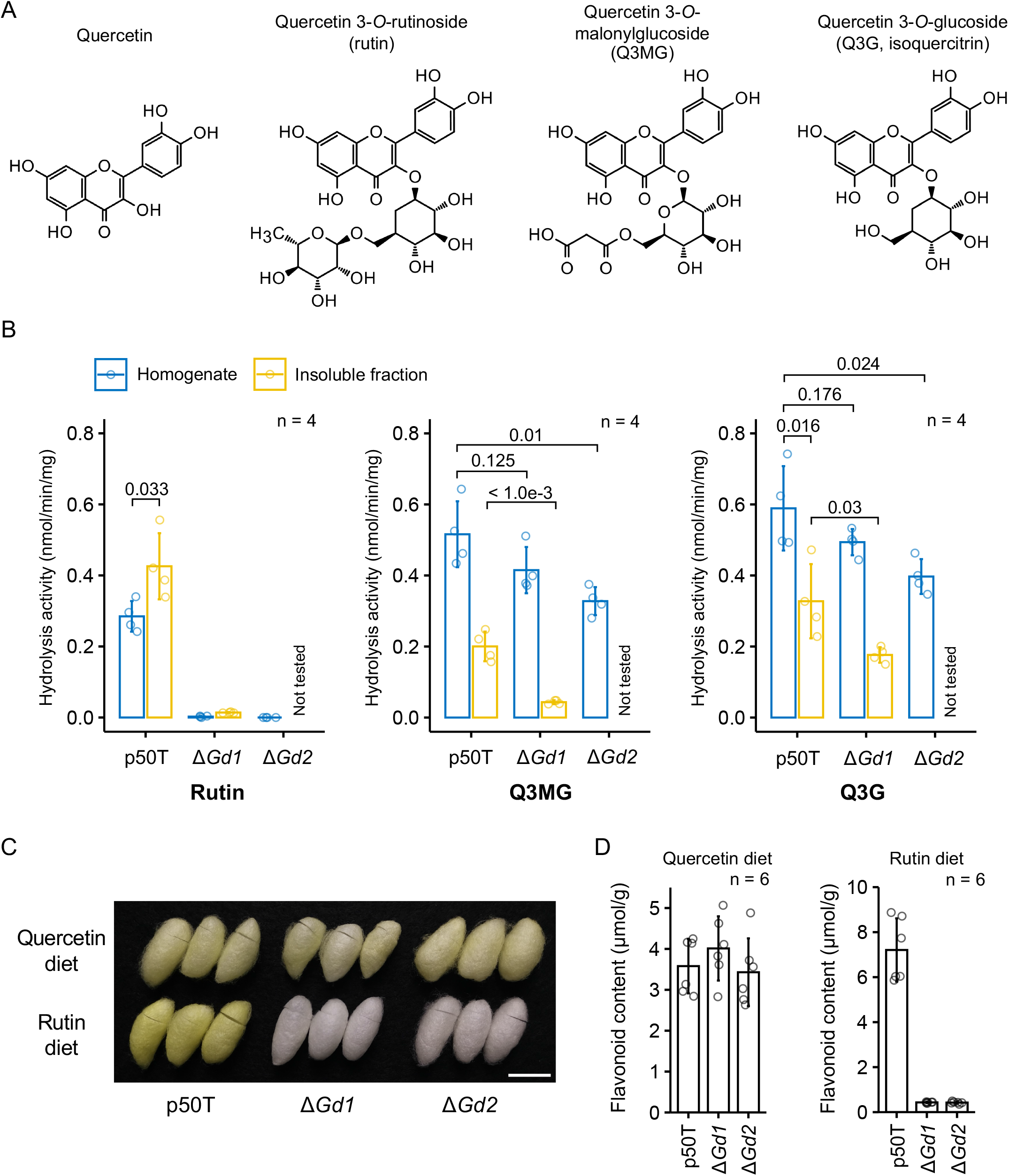
Hydrolytic activity of KWMTBOMO12227 on the three major quercetin glycosides in mulberry leaf. (A) Structural formulae of quercetin and the three major quercetin glycosides present in mulberry leaf. (B) Hydrolytic activity of the midgut on the quercetin glycosides. Data are means ± SD. The values above the graph are *p*-values calculated by a two-tailed Student’s *t*-test. The samples used were all fifth-day final instar female larvae reared on a commercial artificial diet. (C) Photograph of representative cocoons of the p50T strain and the mutants reared on semi-synthetic diets containing quercetin or rutin. Bar = 20 mm. (D) Total flavonoid content in the cocoon of the p50T strain and the mutants reared on semi-synthetic diets containing quercetin or rutin. Data are means ± SD.

The homogenate of the p50T midgut exhibited hydrolytic activity against all three types of quercetin glycoside (Fig 3B). However, the activities were decreased in the knockout mutants, indicating that KWMTBOMO12227 is involved in the hydrolysis of all three quercetin glycosides. Notably, *KWMTBOMO12227* knockout completely abolished the hydrolytic activity against rutin, indicating that KWMTBOMO12227 is the only glycosidase with hydrolytic activity against rutin. The hydrolytic activity of rutin was stronger in the insoluble fraction than in the homogenate, supporting our assertion that KWMTBOMO12227 is a membrane-anchored glycosidase. The reduction in the hydrolytic activity of the other two glycosides, Q3MG and Q3G, was only partial, suggesting that the silkworm has other glycosidases with hydrolytic activities against those molecules (Fig 3B). The reduction in the hydrolytic activity of midgut homogenates against Q3MG and Q3G by knocking out *KWMTBOMO12227* was 0.6 to 0.8-fold and 0.7 to 0.8-fold, respectively. The reduction in the hydrolytic activity of the insoluble fraction against the two quercetin glycosides was 0.2 and 0.5-fold, respectively, which were greater than those of the homogenate. These results suggest that KWMTBOMO12227 is an important protein for the hydrolysis of quercetin glycosides in the silkworm lumen. According to the characteristics of the protein sequence and enzymatic activity of KWMTBOMO12227, we named the identified gene *LQGH1*.

In 1972, Fujimoto and Hayashiya reported that the domestic silkworm accumulates flavonoids in its cocoon when reared on artificial diets containing isolated quercetin or rutin but not when reared on mulberry leaves[29]. To investigate whether the uptake of rutin-derived quercetin is dependent on deglycosylation by LQGH1, we reared insects on semi-synthetic diets supplemented with rutin or quercetin but without mulberry leaf powder and measured the flavonoid content in the cocoons. The p50T strain accumulated flavonoids in its cocoon irrespective of diet (Fig 3C, 3D, and S3 Table). Although the *LQGH1*-knockout mutants accumulated the same amount of flavonoids as did p50T when reared on the quercetin diet, they accumulated only 6% of that accumulated by p50T when reared on the rutin diet (Fig 3C and 3D). These results are consistent with the report by Fujimoto and Hayashiya, and indicate that the uptake of quercetin from rutin is strongly dependent on deglycosylation by *LQGH1*.

### Defective structural mutations of LQGH1 were broadly disseminated in white-cocoon domestic silkworm strains

Although accumulation of carotenoids or flavonoids in the cocoon produces a variety of cocoon colors, it is white-cocoon strains that are the most popular in commercial use. Earlier in the present study, we found that defective mutation of *LQGH1* resulted in impaired quercetin absorption, which made the color of the cocoon closer to white (Fig 2B). Interestingly, the defective mutation of *LQGH1* did not impair the growth of the silkworm, but rather improved silk yield (S5 Fig). The implication here is that this beneficial mutation was selected for in the establishment of white-cocoon domestic silkworm strains.

To examine our hypothesis, we looked at the frequency and distribution of the defective mutation of *LQGH1* in a collection of *B. mori* strains. We first determined the details of the mutation introduced in J01. Previously, we reported a genome assembly and gene model for J01[30]. A BLASTp search of our J01 gene model identified predicted gene *BMN13127* as the corresponding gene of *LQGH1* in J01. Comparing the genomic regions of *LQGH1* in the two strains, an insertion of 3997 bp was found in exon 5 of J01 *LQGH1*. Syntenic dot-plot analysis of the genomic sequences of exons 5, 6 and the intronic region between them (intron 5) of *LQGH1* of the two strains revealed that the inserted region contains the inverted sequence of intron 5 of p50T (Fig 4A). The inverted intron sequence was present at two sites within intron 5 of J01 *LQGH1*. In the genomic PCR amplifying the region on exon 5, including the insertion, an amplified product from the J01 genomic DNA was approximately 4000-bp longer than that from the p50T genomic DNA (Fig 4B). Due to the insertion, the predicted protein sequence of J01 LQGH1 lacked 31 amino acid residues in the middle compared to that of p50T LQGH1 (S6 Fig). The deletion included two of nine residues with >99% conservation across 101 homologous proteins from five Macroheterocera species, and all of seven residues that harbored strongly similar physicochemical properties across the proteins (S7 Fig). These observations strongly suggest that the deletion is mainly responsible for the observed difference in flavonoid content in the cocoon.

**Fig 4.**
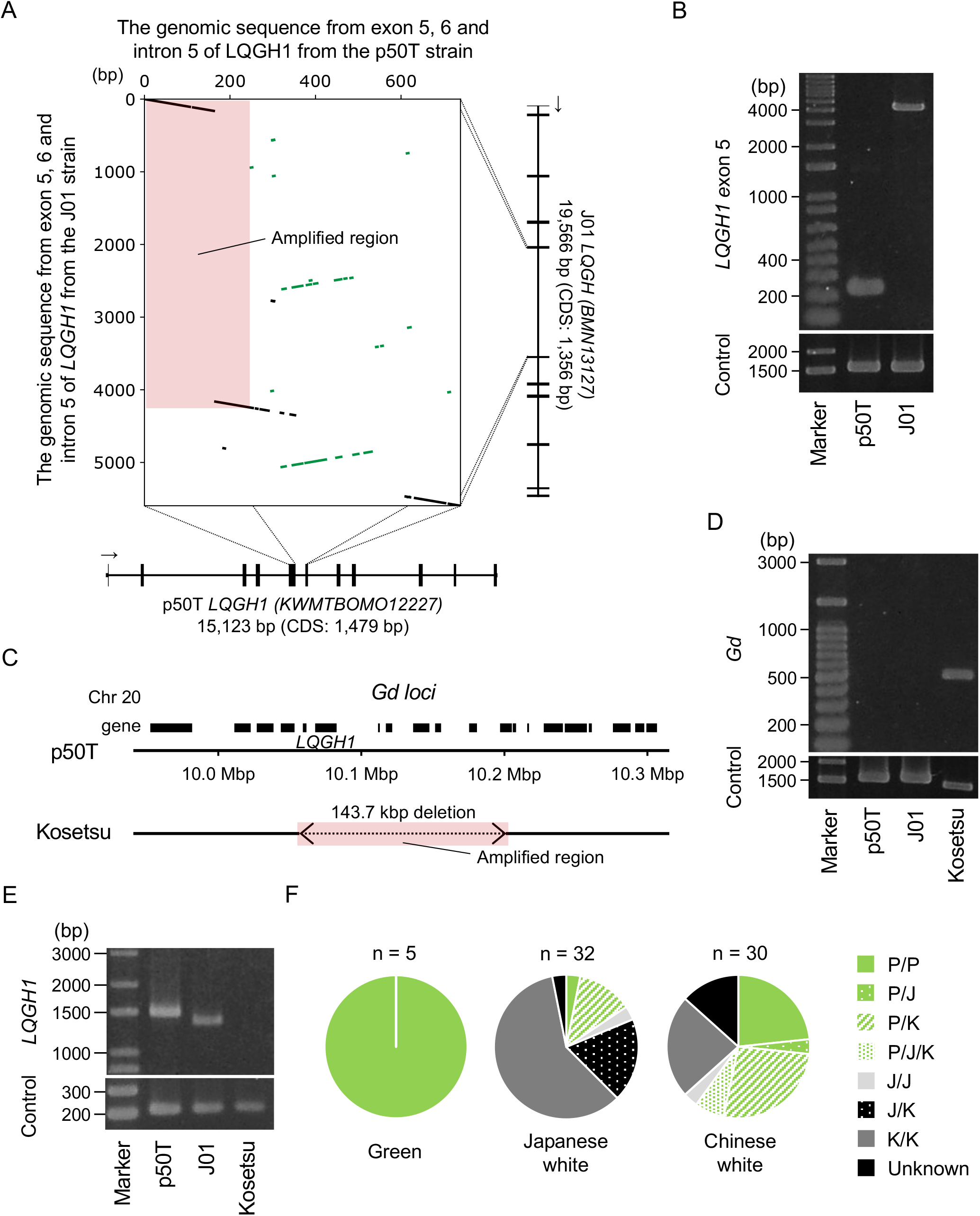
Conserved structural mutations of *LQGH1* in the domestic silkworm population. (A) Dot-plot analysis of the genomic region of exons 5, 6 and the intronic region between them (intron 5) of *LQGH1* from the p50T and J01 strains. (B) PCR identifying the J01-type insertion. The amplified region is indicated by the colored area in (A). The primer set amplifying the genomic region from exons 1 and 2 of *KWMTBOMO14639* (*rp49*) was used as the control (predicted fragment length = 1548 bp). (C) Schematic illustration comparing the *Gd* loci of the p50T and Kosetsu strains. Black boxes indicate the modeled genes. (D) PCR identifying the Kosetsu-type large deletion. The amplified region is indicated by the colored area in (C). (E) RT-PCR of the *LQGH1* full-length open reading frame by using cDNA libraries constructed using the midguts of sixth-day final instar larvae. The primer set targeting exons 1 and 2 of *KWMTBOMO14639* (*rp49*) was used as the control (predicted fragment length = 213 bp). (F) Distribution of the haplotypes of *LQGH1* in Japanese and Chinese local silkworm strains. P/P, only the p50T genotype was detected; P/J/K, p50T, J01, and Kosetsu genotypes were detected; J/J, only the J01 genotype was detected; J/K, both J01 and Kosetsu genotypes were detected; K/K, only the Kosetsu genotype was detected; U, unknown, nothing detected. Grayscale colors indicate pairs of loss-of-function haplotypes.

Next, we examined the presence of the J01-type 4-kbp insertion in 67 Japanese and Chinese local strains by PCR using tested primers (S8 Fig and S4 Table). All five green-cocoon strains lacked the insertion. Of the 32 Japanese local white-cocoon strains, the insertion was absent in five, present in seven, and no amplification product was observed in 20. Of the 30 Chinese local white-cocoon strains, the insertion was absent in 15, present in three (p50T-type products were also observed in these strains), and no amplification product was observed in 11. The lack of an amplification product suggested strains with another dysfunctional mutation. To determine the mutation, we assembled the genomic sequence of one of the strains without an observed amplification product, Kosetsu, using the Illumina whole-genome sequencing data. By comparison against the genomic sequence of p50T, a 143.7-kbp deletion including *LQGH1* was found in the Kosetsu *Gd* locus (Fig 4C). When the region including the deletion was amplified by PCR using p50T, J01, and Kosetsu genomic DNA, an observable amplification product was synthesized only from the Kosetsu sample (Fig 4D). In addition, no observable amplification product was synthesized from the Kosetsu sample by reverse-transcription PCR amplifying the full-length open reading frame of *LQGH1* using cDNA from the midgut of sixth-day final instar larvae of the three strains (Fig 4E). The J01-derived product was shorter in size and had a weaker signal compared to that from p50T. These results are consistent with the genetic differences of *Gd* among the three strains predicted from the assembled genomic sequences. Integrating the genotyping results using primer sets to identify this large deletion, we successfully genotyped the 62 strains. The five strains with no observable amplification product in the PCR have possibly undergone further genome sequence changes by unequal crossing-over after acquiring the Kosetsu-type large deletion (S8 Fig).

To summarize the genomic PCR analysis, three haplotypes of *LQGH1* were found in the population: p50T-type (P), J01-type (J, harboring a 4-kbp insertion in exon 5), and Kosetsu-type (K, harboring a large deletion eliminating *LQGH1*). The proportions of strains homozygous for the functional haplotype (P), heterozygous for the functional and a dysfunctional (J or K) haplotype, and homozygous for a dysfunctional haplotype were 3%, 13%, and 84%, respectively, in the Japanese local white-cocoon strains, and 23%, 37%, and 40%, respectively, in the Chinese local white-cocoon strains (Fig 4F).

## Discussion

### LQGH1 mediates quercetin uptake in *Bombyx mori*

Here, we identified LQGH1 as a glycosidase that initiates quercetin metabolism in the domestic silkworm. LQGH1 is expressed specifically in the midgut and is indicated to be localized on the apical surface of the epithelial cells with a transmembrane domain at the N-terminal (S2 and S4 Fig). Knockout of *LQGH1* reduced total flavonoid content in the midgut, hemolymph, and silk glands, indicating that the gene is involved in quercetin uptake into the midgut (Fig 2C–2F). The hydrolytic activity of the midgut tissue on rutin, Q3MG, and Q3G was reduced in *LQGH1-*knockout mutants (Fig 3B), indicating that LQGH1 hydrolyses all three major quercetin glycosides present in mulberry leaf. The greater rutin hydrolytic activity in the midgut insoluble fraction than in the homogenate supported that LQGH1 is a membrane-anchored protein (Fig 3B). The amount of flavonoids contained in the cocoon of insects reared with a diet supplemented with rutin was greatly reduced in the absence of functional LQGH1 (Fig 3C and 3D), suggesting that deglycosylation of quercetin glycosides by LQGH1 is a critical step in quercetin uptake. However, the present functional analyses were limited to genetic analyses using genome editing. Further biochemical and/or immunological studies will be needed to clarify the enzymatic characteristics and biological role of LQGH1.

While revealing a major role for LQGH1 in quercetin uptake in the silkworm, this study also suggests the presence of LQGH1-independent quercetin glycoside digestion pathways. The accumulation of flavonoids in the cocoon and tissues was not completely lost in the *LQGH1*-knockout mutants (Fig 2F). In addition, the residual hydrolytic activity of the insoluble material fraction of the midgut on Q3MG and Q3G implies the presence of other glycosidases mediating quercetin uptake (Fig 3B). Although we were unable to successfully obtain lineages in which *KWMTBOMO12222* and *KWMTBOMO12236* were knocked out, their proteins are possibly the identities of these other glycosidases. In addition, the hydrolytic activity of Q3MG and Q3G in the *LQGH1*-knockout mutant was higher in the homogenate than in the insoluble material fraction (Fig 3B), implying that digestion of quercetin glycosides also occurs in the midgut cells by unanchored glycosidases.

### Adaptive significance of quercetin uptake in mulberry silkworm is supported by the fact that LQGH1 has acquired and conserved rutin hydrolytic activity

Our results show that LQGH1 mediates quercetin uptake in the domestic silkworm in a deglycosylation-dependent manner, similar to LPH in mammals. However, our data indicate that *LQGH1* is not an ortholog corresponding to mammalian *LPH*, rather it has evolved through gene duplications during Lepidoptera diversification and has acquired a peculiar function, rutin hydrolytic activity (Fig 3, S10 and S11 Figs, S1 Text). *In vitro*, LPH hydrolyses Q3MG and Q3G, but not rutin [24,26,31]. Rutin hydrolysis in mammals is dependent on the intestinal microbiota[32]. It also depends on the gut microbiota in the honey bee (*Apis mellifera*)[33]. Rutin glycosidases have previously been identified in bacteria, fungi, and plants [34,35,36], but to our knowledge, not in animals. Rutin is a common plant flavonoid, accounting for 14%–46% of flavonol glycosides in mulberry leaf[17]. The accumulation of flavonoids in the cocoon of *LQGH1*-knockout mutants reared on a rutin-containing diet was severely restricted, indicating the absence of rutin digestion pathways other than deglycosylation by LQGH1 (Fig 3C and 3D). Therefore, the acquisition of rutin hydrolytic activity should result in a considerable change in quercetin intake. In addition, the phylogenetic tree of the lepidopteran proteins indicated that frequent gene duplication events in the *Gd* glycosidase family have occurred (S11 Fig). In the inference of gene duplication events by Orthofinder[37], the number of events in the LQGH1-included group was the most frequent among 55 ortholog groups including 78 silkworm proteins annotated as glycosidases (S5 and S6 Table). While the genomic instability of the *Gd* locus had possibly provided genetic materials to evolve *LQGH1* by gene duplication events, it can be inferred to have repeatedly exposed *LQGH1* to a crisis of loss by unequal crossing-over. However, *LQGH1* remained conserved until its defective mutation was selected in breeding. This emphasizes the significance of quercetin bioavailability brought about by *LQGH1*; our study supports that the suggested positive properties of quercetin glycosides, their antioxidant[15], ultraviolet light-protective[14], and antibacterial functions[16], improve the fitness of the wild silkworm in the natural environment. Although, it should be noted that excessive quercetin uptake is still toxic even for the domestic silkworm[38], meaning that the mulberry silkworms should have a mechanism to maintain appropriate levels of quercetin. The fluorescence in the posterior region of the midgut observed in the present study implied accumulation of quercetin glucosides with glycosylation at the 5-*O* position and the reloading of excessive quercetin glucosides from the hemolymph to excrete them into the midgut lumen (Fig 2C).

### *LQGH1* loss contributed to cocoon improvement

Through breeding, many strains of the domestic silkworm have been established that exhibit a wide variety of cocoon colors depending on their flavonoid and carotenoid content, but commercially white cocoons tend to be preferred in regions such as Japan and China, especially since the 20th century. In a previous study that conducted a worldwide genetic analysis of silkworm strains, 91 of the 121 strains examined were white-cocoon strains[39]. The defective mutation in *LQGH1*, which manifests as reduced cocoon color (Fig 2B), may have contributed to the establishment of these white-cocoon silkworm strains. Of the white-cocoon strains examined in the present study, the proportion with homozygous defective haplotypes of *LQGH1* (haplotype J or K) was only 62% (Fig 4F, S8 Fig, and S4 Table). In 24% of the strains, a mixture of defective and functional haplotypes (haplotype P) was present. These are not surprising results, considering that not all white-cocoon strains harbor loss-of-function mutations of the sugar transporters on the *Gn* locus, which is more effective than *Gd* with respect to cocoon discoloration[18]. Even if visually determined as white, the actual color tone and therefore the flavonoid content of the cocoon varies from strain to strain[40]. In addition to *Gd*, *Gb*, and *Gn*, within each of which a gene involved in the quercetin metabolism has been identified, several loci associated with cocoon greenness have been reported, including *Gc* [21,22], *Green cocoon* (*Grc*), *Green egg shell* (*Gre*), and *Yellow fluorescent* (*Yf*)[21]. “White cocoon” is presumed to be expressed by additive cocoon color discoloration by a combination of loss-of-function haplotypes of these loci.

Although loss of *LQGH1* is not necessarily essential for cocoon whiteness, given the high conservation rate of loss-of-function haplotypes and the fact that knockout of the gene reduced the cocoon total flavonoid content by less than half (Fig 2F, Fig 4F), the gene still makes an important contribution to cocoon discoloration in the domestic silkworm. Interestingly, the knockout of *LQGH1* increased the cocoon weight (S5 Fig). Because endoreduplication is an important step in silkworm silk gland development[41], interference with the replication step by quercetin interaction with DNA may explain the relationship between *LQGH1* and silk yield[42]. Although it will be difficult to confirm this hypothesis because cocoon weight is a complex quantitative trait dependent on silkworm physiology and behavior, elucidation of the relationship between quercetin and silk yield may provide clues to further improve silkworm protein productivity.

## Materials and Methods

### Insect materials

All domestic silkworm strains used in the present study were maintained at the National Agriculture and Food Research Organization (NARO, Japan). p50T, the strain used for the functional analysis of *LQGH1*, is almost genetically identical to p50, because the strain is a descendant strain of p50 that had undergone repeated passage using a single pair to improve homozygosity for referential use. The silkworms were reared on a commercial artificial diet (SilkMate PM; Nosan, Kanagawa, Japan) or fresh mulberry leaves under a controlled environment (12-h light/dark photoperiod, 25°C). Insects reared on fresh mulberry leaves were used for measurement of the flavonoid contents of the cocoons of p50 and J01 (Fig 1A and 1B), the scoring for QTL analysis (Fig 1C), and measurement of the flavonoid contents of the cocoons of p50T and *LQGH1*-knockout mutants (S3 Fig). All individuals, except those from which genomic DNA and RNA-seq data were derived, were female.

### Flavonoid content measurement

Flavonoids in cocoons were extracted with MeOH–H2O (7:3, V/V) at 60°C for 2 h, after shredding the cocoons into 2–3-mm squares. The extraction was repeated twice for thorough extraction. The flavonoid content in the solution was derived from the absorption at 365 nm, which is highly correlated with flavonoid content (*r* = 0.95)[19]. In the case of flavonoid content as an input for QTL analysis, the raw values of the absorbance were used as relative phenotype scores. The tissues and organs for measuring flavonoid content were sampled from sixth-day final instar larvae reared on the commercial artificial diet, rinsed well with PBS buffer (pH 7.4) (Takara Bio, Shiga, Japan), and stored at −80°C. Immediately after collection, to inhibit melanization, 0.5 M sodium dimethyldithiocarbamate (Fujifilm, Tokyo, Japan) was added to reach 0.5% by volume to the collected hemolymph. Quantification of flavonoid content in the samples was performed according to a previous report[15] with some modifications. In brief, 30 µL of sample was injected into an LC-10ADVP HPLC system equipped with an SPD-10AV photodiode array detector (both Shimadzu Co., Kyoto, Japan) and separated with a Sunfire C18 column (150 × 3.0 mm i.d.; Waters, Massachusetts, USA) at a flow rate of 1.0 mL/min. Spectrum analysis and flavonoid peak assignment were performed with the CLASS-VP HPLC software (Shimadzu Co.). Flavonoids were quantified using quercetin 5, 4′-di-*O*-glucoside as the reference compound.

### QTL analysis

The intercross F2 population of p50T and J01 used in the present study were the same samples as those sequenced by double-digest restriction-associated DNA sequencing and used to generate linkage maps in our previous study[30]. The sequencing data of 102 female F2 individuals are available in the Sequence Read Archive under the BioProject accession number PRJDB13956. The raw values of absorbance, indicating the relative cocoon flavonoid content for each individual, are summarized in S1 Table. The method for genotyping, marker selection, and construction of linkage maps is described in detail in our previous report[30]. Briefly, OneMap v2.8.2[43] was used to construct linkage maps with an LOD score of 3, with the genotype of the F2 population output from the script “ref_map.pl” in STACKS v1.48[44] as the input. The p50T genome assembly was used as the reference for read mapping[23]. The linkage map used consisted of 1038 makers, covered a total genetic length of 945.36 cM, and had an average marker density of 0.936 cM. QTL analysis was performed using the R/qtl v1.46.2 package[45], with the relative flavonoid content scores in the cocoons and the genetic position of markers linked to the physical position on the p50T genomic sequence as inputs. Using the “calc.genoprob” function in R/qtl, the probabilities of the true underlying genotypes were calculated with a step size of 1 cM and an assumed genotyping error rate of 0.05. QTL detection was performed by composite interval mapping using the function “cim” with the following parameters: method = “hk”, n.marcovar = 3, window = 10. The LOD significance threshold for detecting QTL was calculated by a permutation test of 1000 trials. Approximate Bayesian 95% credible intervals were calculated using the function “bayesint”. The positions of the nearest markers outside of the boundaries of a confidence interval were defined as the ends of the interval. PVEs and additive effects of the three significant QTLs were estimated using the function “fitqtl” with the parameter: formula = y ∼ Q1 + Q2 + Q3 + Q1 * Q2 * Q3.

### Gene editing

CHOPCHOP v3 was used to identify target sites[46]. Each 20-nucleotide guide sequence was unique on the p50T genomic sequence[23]. The construction of DNA templates for the synthesis of single-guide RNA (sgRNA) was performed according to a previous report[47]. Briefly, the DNA template, which included the T7 promoter, sgRNA target with PAM motif, and tracrRNA-complemental sequence, was synthesized by template-free PCR. A MEGAshortscript T7 Transcription Kit (Thermo Fisher Scientific, Waltham, Massachusetts, U.S.A.) was used for *in vitro* sgRNA transcription and template DNA digestion. A Guide-it IVT RNA Clean-Up Kit (Takara Bio) was used to clean up the sgRNA solution. Alt-RS.p. HiFi Cas9 Nuclease V3 solution (Integrated DNA Technologies, Coralville, Iowa, U.S.A.) and the sgRNA were mixed with distilled water to final concentrations of 500 ng/μL and 50 ng/μL, respectively. The Cas9 nuclease and sgRNA mixed solution was incubated for 1 h at room temperature to allow the ribonucleoprotein to form. The ribonucleoprotein solution was microinjected into non-diapause eggs of the p50T strain within 8 h after oviposition, in accordance with a previous report[48]. Non-diapause treatment was performed by incubating the eggs under 17°C short-day conditions (12-h light/dark photoperiod) until hatching. Adult moths of G0 individuals (injected generation) were crossed with wild-type moths. High-resolution melting (HRM) analysis using genomic DNA extracted from molt shells of the G1 generations in cocoons identified individuals harboring identical heterozygous mutant haplotypes. PCR for HRM analysis was performed by using a KAPA HRM Fast PCR Kit (Roche, Basel, Switzerland) on a LightCycler 96 System (Roche) according to the manufacturer’s instruction. Genomic DNA extraction from molt shells was performed by homogenizing the shells in a typical SDS-based lysis buffer followed by isopropanol precipitation. By the same method, G2 individuals with a homozygous mutant haplotype were identified and crossed, establishing the knockout lines. PCR products including the target site were sequenced using a BigDye Terminator v3.1 Cycle Sequencing Kit and an Applied Biosystems 3130xl Genetic Analyzer (Thermo Fisher Scientific) to confirm the mutation. The primer sequences used are listed in S7 Table.

### Photographing of dissected samples

Sixth-day fifth instar larvae reared on the artificial diet were used. Photographs were collected with a DP74 camera (Olympus, Tokyo, Japan) attached to an SZX16 microscope (Olympus). Immediately after collection, to inhibit melanization, 0.5 M sodium dimethyldithiocarbamate (Fujifilm) was added to reach 0.5% by volume to the collected hemolymph. Silk glands and whole-body samples from which silk glands had been removed were rinsed well with PBS buffer (pH 7.4) (Takara Bio) before imaging to remove fluorescent hemolymph. A fluorescence light source U-HGLGPS (Olympus) and ultraviolet light filter set (excitation: 330–385 nm, emission: 420 nm longpass) (Olympus) were used for imaging under ultraviolet light irradiation. Because of the loss of resolution and optical noise when the field of view was made large enough to fit the whole body, the whole-body samples were photographed in segments and then the pictures were combined. In the photographs shown in Fig 2, no post-editing of any kind, other than cropping, was performed.

### Isoform identification and expressional analysis

RNA-seq data from the midgut and other organs of third-day fifth instar larvae of p50T and J01 were previously published by our research group [30,49]. The sequencing reads were trimmed by fastp v0.20.0[50] with the following parameters: −q 20 −n 5 −l 100. The mapping of the reads to the p50T genome assembly and transcript isoform identification were conducted using HISAT2 v2.1.0[51] and SrtingTie2 v2.2.0[52] with the default parameters. Salmon v1.5.2[53] was used to calculate transcripts per kilobase million scores. The modeled transcript sequence of p50T was used as the reference[23].

### Phylogenetic analysis

To collect protein sequences for constructing the phylogenetic tree of glycosidases of mammals and insects (S10 Fig), the related proteins were collected by BLASTp search using the predicted protein sequence of *KWMTBOMO12227* of the p50T strain gene model as the query sequence[23]. Genes with e-values less than 1E-50 were used for the analysis. The KEGG[54] BLAST search service (https://www.genome.jp/tools/blast/) was used for picking related proteins in *Rattus norvegicus*, *Drosophila melanogaster*, *Tribolium castaneum*, and *Apis mellifera*. The query and these sequences were aligned using Clustal Omega v1.2.4[55]. Tree construction by the neighbor-joining method was performed using MEGA 11[56]. The number of bootstrap trials was set to 100. The phylogenetic tree of lepidopteran orthologs of the *Gd* glycosidases was constructed by the maximum likelihood method using RAxML-NG v1.1[57] with the following parameters: --bs-trees 100, --model LG+I+G4 (S11 Fig A). The best-fit phylogenetic analysis model was suggested using ModelTest-NG v0.1.7[58]. Gene models of 12 lepidopteran insects available in InsectBase 2.0[59] (http://v2.insect-genome.com/) were used (*Papilio Xuthus*, *Megathymus ursus*, *Lycaena phlaeas*, *Limenitis camilla*, *Chilo suppressalis*, *Biston betularia*, *Pheosia gnoma*, *Dendrolimus punctatus*, *Bombyx mori*, *Bombyx mandarina*, *Antheraea yamamai*, *Deilephila porcellus*); each of the models were predicted based on high-quality genome sequences using long-read sequencing, and their BUSCO[60] scores were all above 94%. Rat LPH (NCBI RefSeq accession: NP_446293) and the total of 90 protein sequences of the lepidopteran species determined by Orthofinder[37] v2.5.2 to belong to the same ortholog group as any of the nine *Gd* glycosidases were used in the analysis.

### Assay for hydrolytic activity of the midgut

Midguts of fifth-day final instar larvae reared on the commercial artificial diet were collected, rinsed well with PBS buffer (pH 7.4) (Takara Bio), and immediately stored at −80°C. Three times the weight of 10 mM phosphate/1 mM PMSF buffer (pH 7.5) as the samples was added and they were homogenized. Centrifugation at 20,000*g* for 20 min, collection of the precipitate fraction, and washing with the buffer were repeated twice for the homogenate sample to obtain an insoluble fraction containing cell membranes. Hydrolysis was performed in a 20 mM phosphoric acid (pH 5.5) solution with quercetin glycosides at a concentration of 2 mM for 4 h at 37°C. The reaction was stopped by adding three volumes (V/V) of methanol. Control incubations in which the substrates were omitted were also run. After centrifugation at 20,000*g* for 10 min, the quercetin content of the supernatant was quantified with the HPLC system as described in the section “Flavonoid content measurement.” Quercetin, rutin, and Q3G were obtained from Extrasynthese (Lyon, France). Q3MG was obtained from Merck (Darmstadt, Germany). Hydrolytic activity toward the quercetin glycosides is expressed as nanomoles of quercetin produced per min per mg protein. The protein concentration in the enzyme preparations was measured with a commercial assay kit (Coomassie Plus, Thermo Fisher Scientific).

### Dietary administration of quercetin or rutin

Silkworm larvae were reared on the commercial diet from hatching to the third ecdysis, then the fourth instar larvae were fed with a diet containing 25% mulberry leaf powder[61]. The newly molted fifth instar larvae were reared on semi-synthetic diets supplemented with quercetin or rutin, which did not contain mulberry leaf powder (S3 Table). Flavonoids added to the diets were approximately equimolar (0.1 mmol/100-g dry diet). Chlorogenic acid was added to the diets as a feeding stimulant. Soybean meal (Soya flour FT; Nisshin OilliO Group,. Ltd., Tokyo, Japan) was washed twice with five volumes of 90% (V/V) ethanol to remove deterrent substances and allowed to dry naturally before use in the diets.

### Genomic comparison of the *Gd* locus

Dot-plot analysis was performed using FlexiDot v1.06[62] with the following parameters: -p 1 -f 1 -A 2 -E 100. The illustrated exon-intron structure of J01 *LQGH1* (*BMN13127*) was modified from the original model (Fig 4A); the incorrect prediction of exon 1 was modified according to *KWMTBOMO12227*, and the J01 transcript sequence was obtained by using the original RNA-seq data[30]. DNA extraction of the domestic silkworm strain Kosetsu from the whole body of a male larva was performed by a typical phenol-chloroform-based method. Whole genomic sequencing was performed using an Illumina NovaSeq 6000 Sequencing System (Illumina Inc., San Diego, USA) via an outsourcing service (Macrogen Japan Corp., Kyoto, Japan). The sequencing generated 50.4 Gb of paired-end reads with a length of 151 bp. Genome assembly of the Kosetsu strain was performed using MaSuRCA v4.0.9[63] with pseudomolecule sequences of chromosomes 15, 20, and 27 of p50T[23], on which the QTLs are present, as the assembly reference. The parameter defining 20 times genomic size “JF_SIZE” was set to 9,000,000,000. The input genomic sequencing data of the Kosetsu strain was trimmed by fastp v0.20.0[46] with the following parameters before assembly: -q 20 -n 5 -l 100.

### PCR-based genotyping

The genomic DNA of each strain was extracted from 10 pairs of silk glands of final instar larvae by a typical phenol-chloroform-based method. The final concentration of genomic DNA was conditioned to 0.5 ng/μL. KOD One PCR Master Mix-Blue-(Toyobo, Osaka, Japan) was used for PCR. The times and temperatures of preincubation, denaturation, annealing, and extension were set to 30 s at 95°C, 10 s at 98°C, 10 s at 60°C, and 45 s at 68°C, respectively, for 32 cycles. The primer sequences used are listed in S7 Table.

### Reverse-transcription PCR

Total RNA was extracted from the midgut of sixth-day final instar larvae using ISOGEN (Nippon Gene, Tokyo, Japan) according to the manufacturer’s instructions. Contaminating DNA in the RNA solution was digested using RNase-Free DNase Set (Qiagen, Hilden, Germany). The isolated RNA was purified using an RNeasy Kit (Qiagen). Equal volumes of RNA solutions from five individuals, adjusted to the same concentration, were mixed to obtain a single bulk sample. Reverse transcription was performed using ReverTra Ace qPCR RT Master Mix (Toyobo) according to the manufacturer’s instruction, and the final concentration of the RNA was conditioned to 10 ng/μL. RT-PCR was performed using KAPA HiFi HotStart ReadyMix (Roche) under conditions that resulted in 1% of the concentration of the original cDNA solution. The times and temperatures of preincubation, denaturation, annealing, and extension were set to 180 s at 95°C, 5 s at 98°C, 10 s at 60°C, and 60 s at 72°C, respectively, for 28 cycles. The primer sequences used are listed in S7 Table.

### Detecting gene duplication events

Detecting gene duplication events in lepidopteran *Gd* glysosidases was performed using Orthofinder v2.5.2[37] together with the ortholog-group inference for the phylogenetic analysis. The glycosidase gene was identified in the domestic silkworm by referencing the annotation file of silkworm protein using InterProScan[64] within KAIKObase[65] (https://kaikobase.dna.affrc.go.jp/index.html). Proteins annotated as “Glycoside hydrolase superfamily (IPR017853)” or with the functional annotation “hydrolase activity, acting on glycosyl bonds (GO:0016798)” were considered to be proteins annotated as glycosidases.

### Other informatic tools

Handling sequences: Seqkit v2.2.0[66]; statistical calculations: R v4.0.3 (R Foundation for Statistical Computing, Vienna, Austria); illustrating graphs: ggplot2 v3.4.1[67] and ggpubr v0.6.0 (https://CRAN.R-project.org/package=ggpubr); prediction of hydrophobic transmembrane domains: SOSUI v1.11[68]; illustrating structural formulae: Ketcher v2.8.0 (https://github.com/epam/ketcher); genomic browsing: Integrative Genomics Viewer v2.12.3[69]; phylogenetic tree editing: Interactive Tree Of Life v5[70].

## Supplementary information

Supplementary materials are available at Figshare. doi: <u>10.6084/m9.figshare.23553015</u>

## Data availability

The raw genomic sequencing data of the Kosetsu strain are available in the Sequence Read Archive under the BioProject accession number PRJDB15366.

## Acknowledgements

We thank Ms. Mayuko Sakatoh for her technical assistance; Sae Furuhashi, Toshihiko Misawa, Kaoru Nakamura, Keita Hiruma, Koji Hashimoto, and Eiji Okada for their support in the silkworm rearing; Hiroki Sakai for giving instruction on the CRISPR-Cas9 system construction; and Nobuto Yamada for giving instruction on the microinjection. The computation was partly performed using the supercomputer of the Agriculture, Forestry and Fisheries Research Information Technology Center of the Japanese Ministry of Agriculture, Forestry and Fisheries.

## Author contributions

RW: Conceptualization, Data Curation, Formal Analysis, Investigation, Visualization, Writing – original draft preparation, Writing – review & editing; CH: Conceptualization, Investigation, Writing – review & editing; ST: Resources, Writing – review & editing, Supervision; TI: Conceptualization, Funding Acquisition, Investigation, Project Administration, Resources; SK: Investigation; AJ: Investigation, Writing – review & editing; TT: Resources, Writing – review & editing; KYo: Resources; KYa: Supervision; HS: Funding Acquisition, Supervision

## Description of supplementary materials

**S1 Fig.**
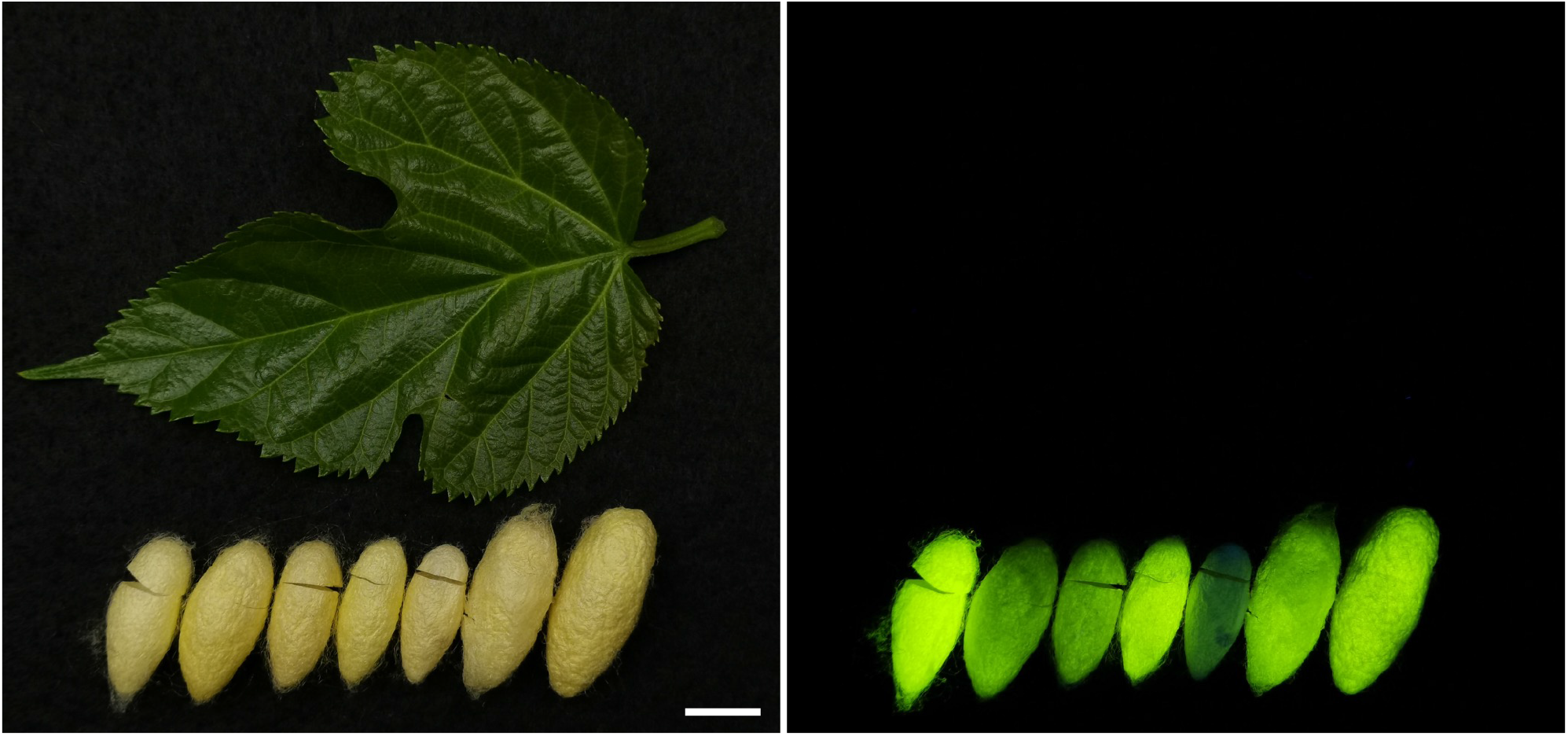
Photographs of representative cocoons of the wild silkworm, *Bombyx mandarina*. A fresh mulberry leaf and cocoons of the wild silkworm in a bright field (left) and those irradiated with ultraviolet light A in a dark field (right). The cocoons under ultraviolet irradiation exhibit fluorescence that is characteristic of quercetin-5-*O*-glucoside and quercetin-5,4′-di-*O*-glucoside, the major quercetin metabolites in the silkworm tissues and cocoon [14,15,28]. The cocoons were collected in June 2023 at Tsukuba, Ibaraki, Japan. Bar = 10 mm.

**S2 Fig.**
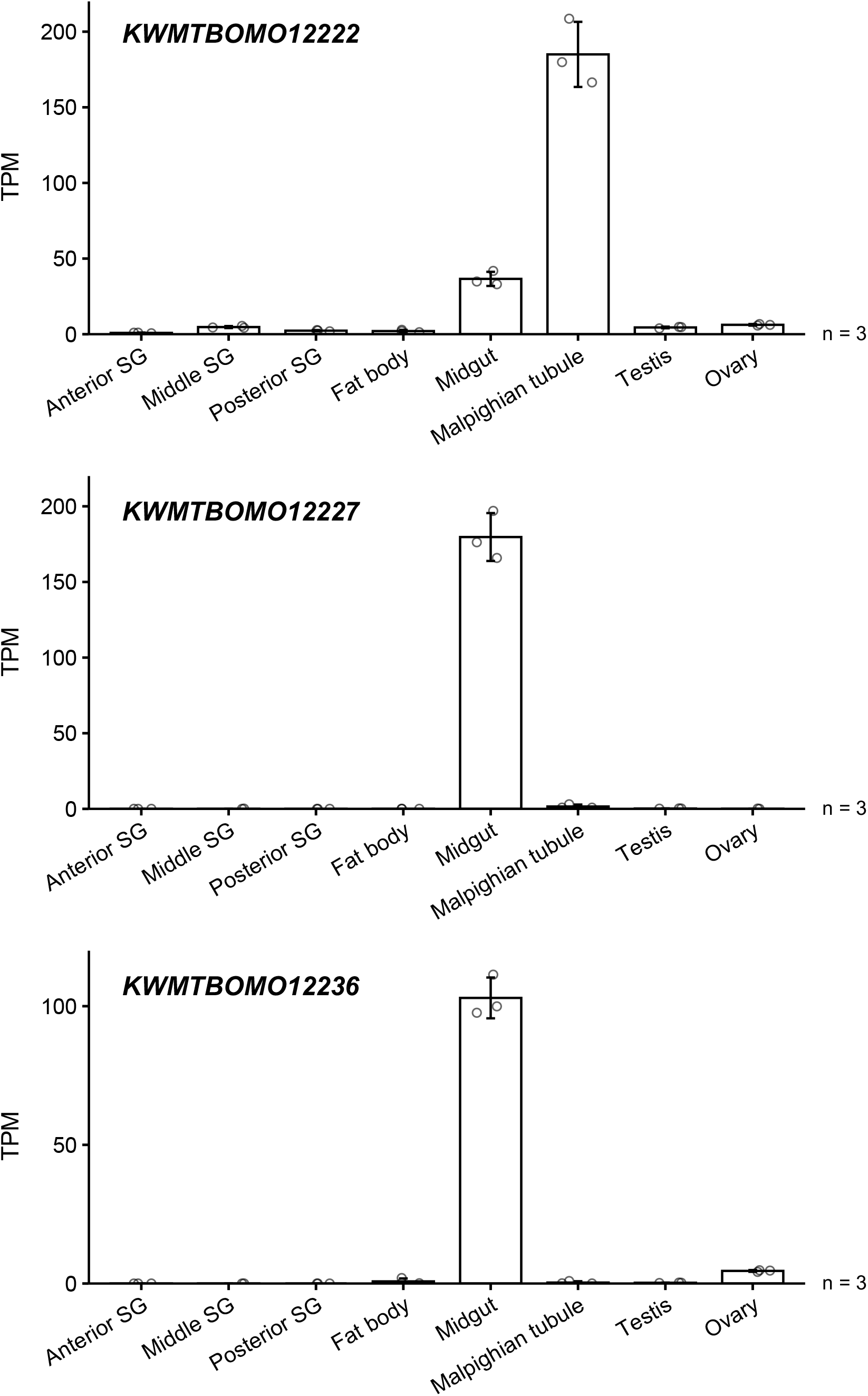
Expression profile of the candidate genes within the *Gd* locus in third-day final instar larva of the p50T strain. All organs and tissues, except for ovaries, were taken from male larvae. Data are means ± SD. SG, silk gland; TPM, transcripts per million.

**S3 Fig.**
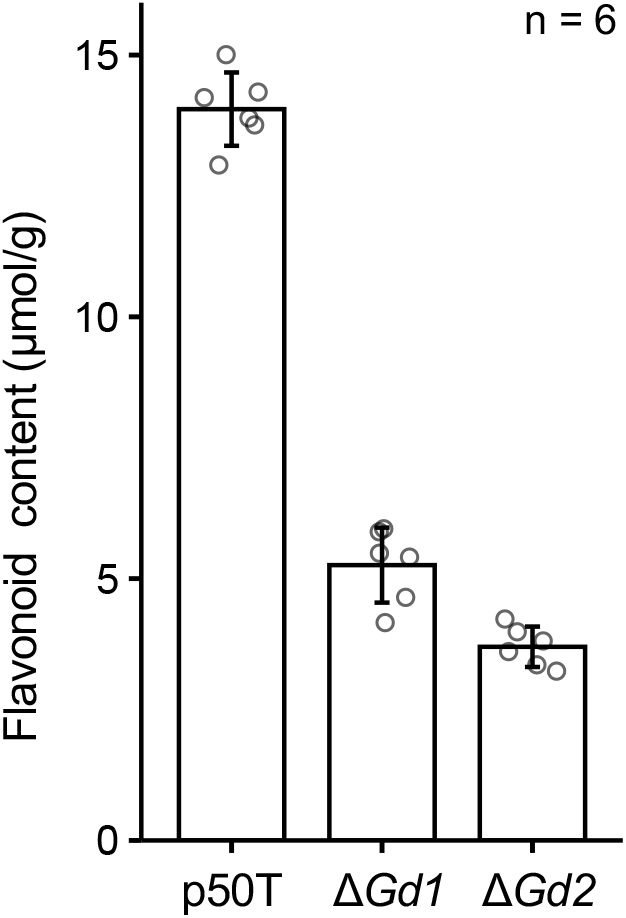
Total flavonoid content in the cocoons of the p50T strain and the *KWMTBOMO12227*-knockout lineages reared on fresh mulberry leaves. Data are means ± SD.

**S4 Fig.**
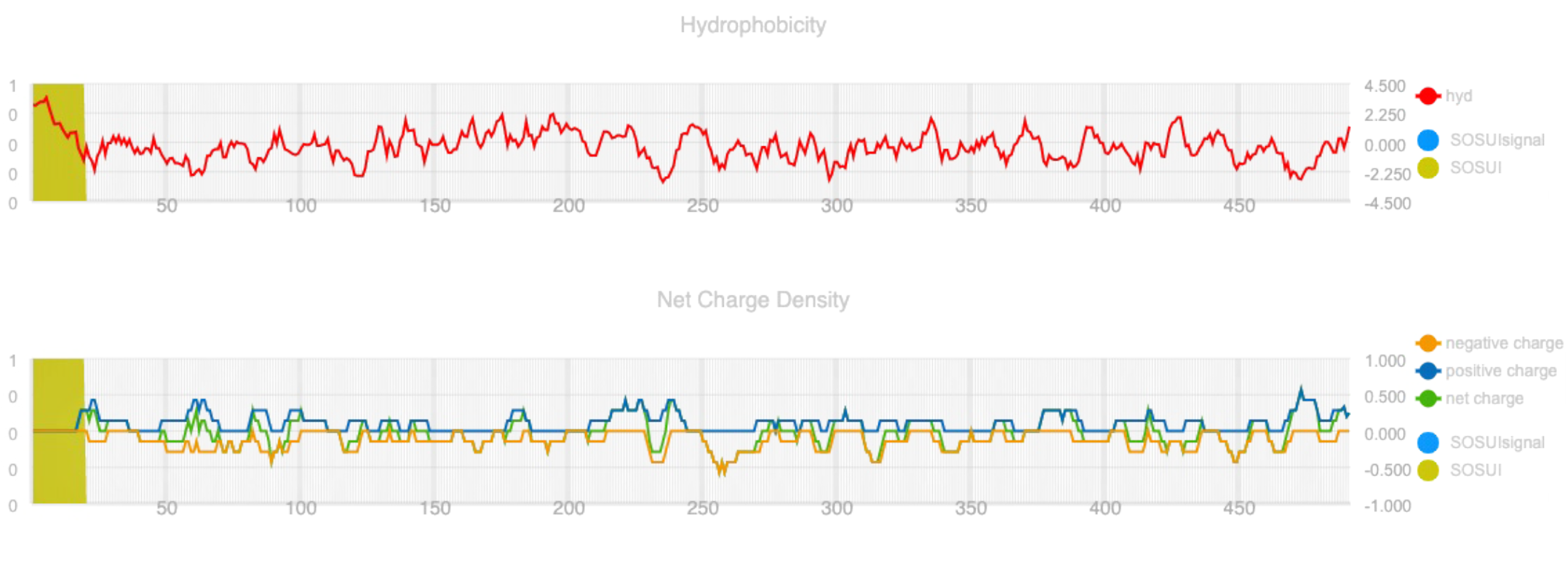
Prediction of a transmembrane domain in the amino acid sequence of KWMTBOMO12227 by using the SOSUI tool.

**S5 Fig.**
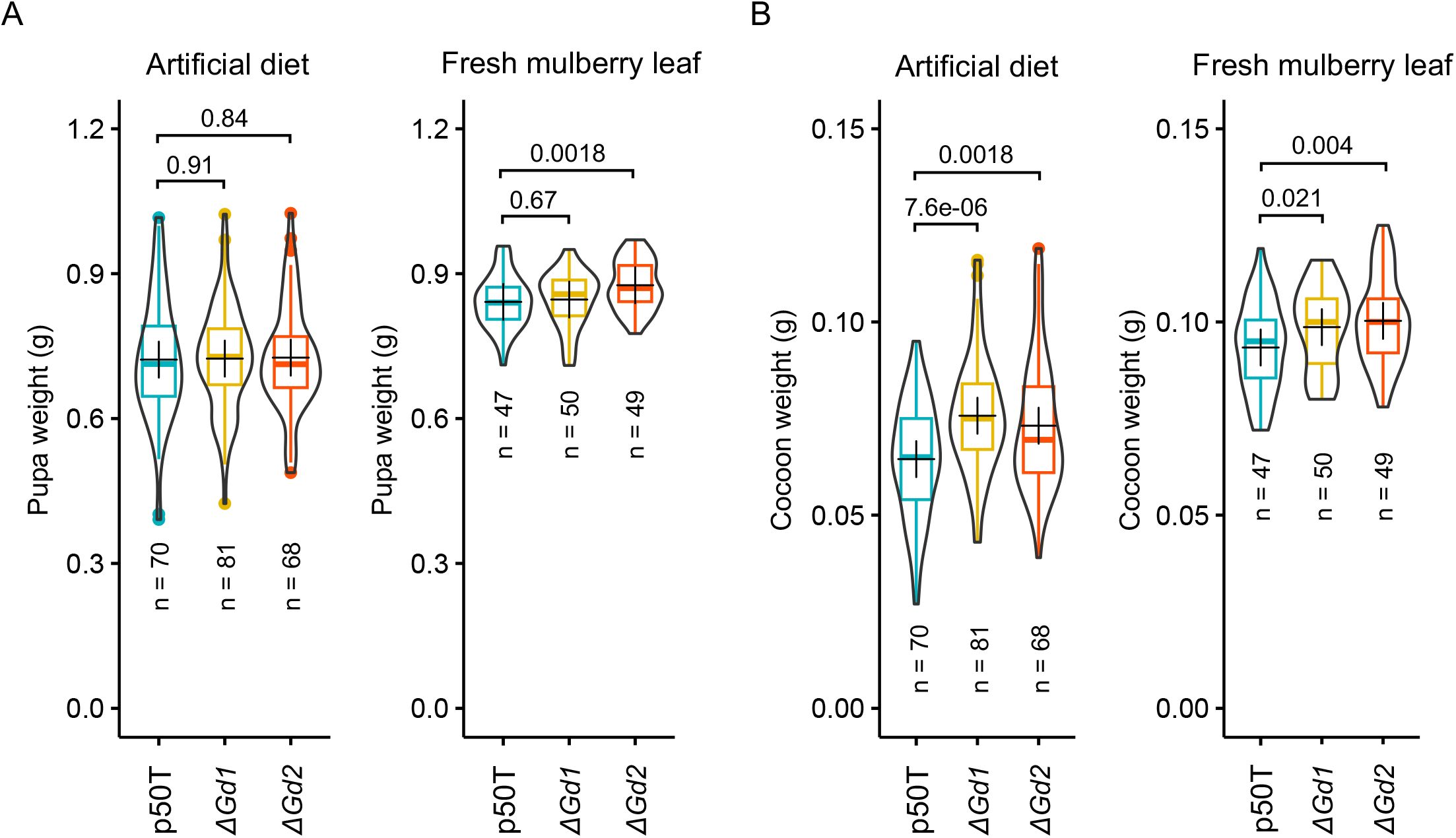
Weight of pupa and cocoon of p50T and the knockout mutants of *LQGH1*. (A) Weight of the pupa. (B) Weight of the cocoon. The values above the graph are *p*-values calculated by Student’s *t*-test. The crosses inside the boxes indicate the means.

**S6 Fig.**
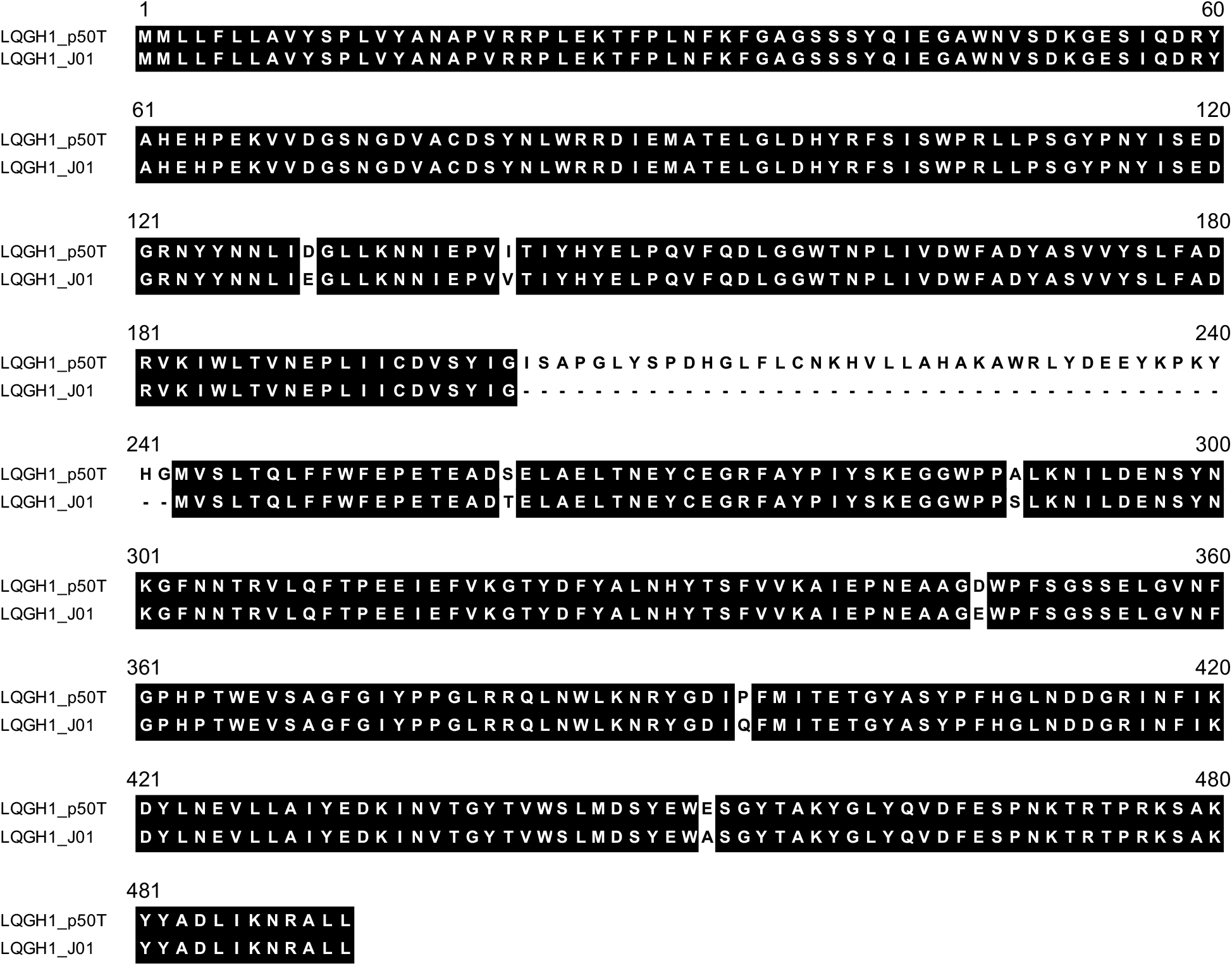
Alignment of the amino acid sequences of LQGH1 from the p50T and J01 strains. Amino acid sequences were aligned using Clustal Omega.

**S7 Fig.**
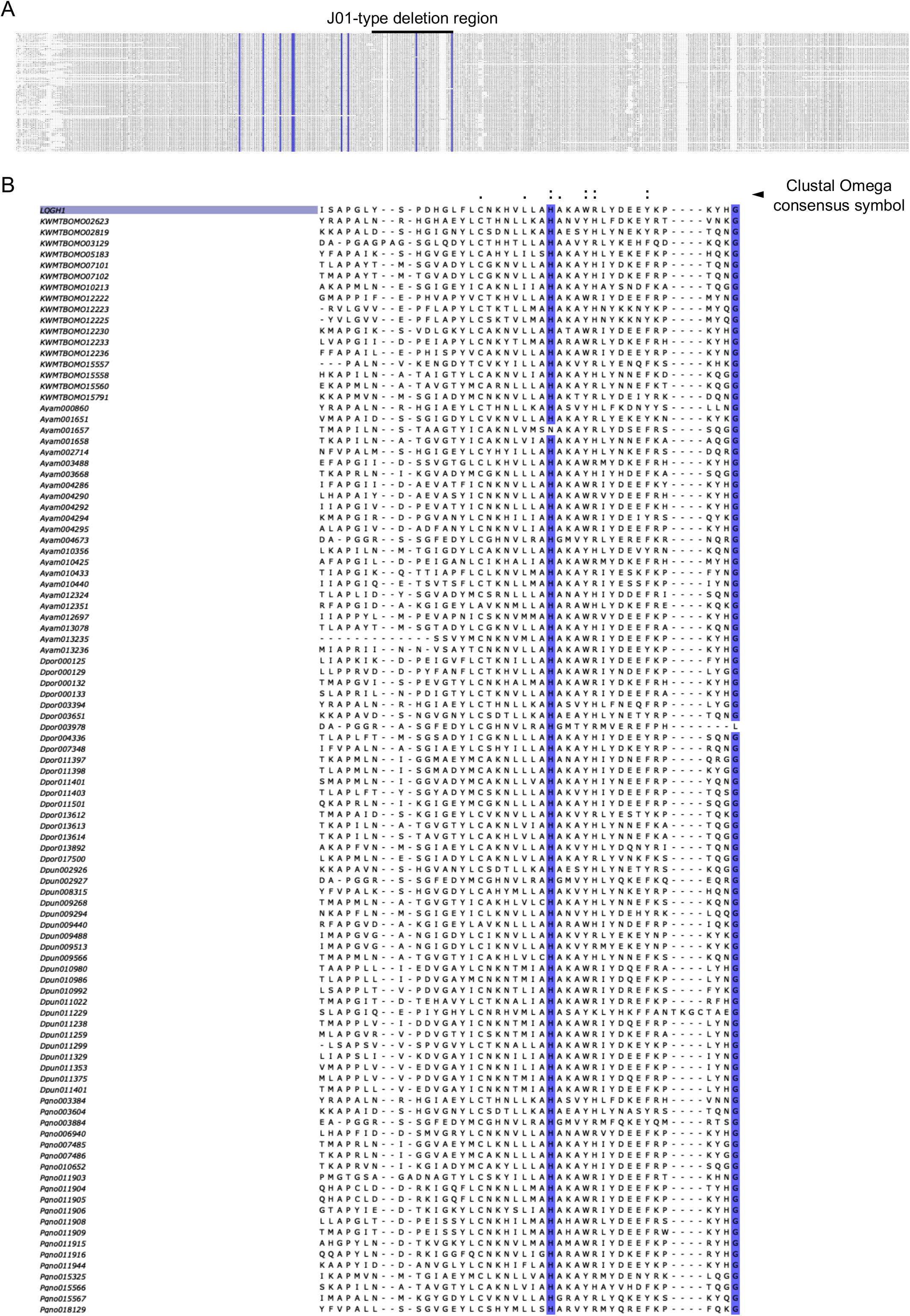
Alignment of LQGH1-like proteins in Macroheterocera species. (A) Whole alignment of 101 LQGH1-like proteins in Macroheterocera species. Amino acid residues identical to LQGH1 at positions with >99% conservation are highlighted by blue. The 100 homologous proteins were collected by a BLASTp search using the amino acid sequence of LQGH1 from each protein sequence data of *Bombyx mori* (KWMT), *Antheraea yamamai* (Ayam), *Deilephila porcellus* (Dpor), *Dendrolimus punctatus* (Dpun) and *Pheosia gnoma* (Pgno) with a cutoff criterion e-value of <1E-100. The sequence data of *B. mori* were obtained from KAIKObase (https://kaikobase.dna.affrc.go.jp/index.html) and those of *A. yamamai*, *D. porcellus* and *P. gnoma* were obtained from InsectBase 2.0 (http://v2.insect-genome.com/). The sequences were aligned using Clustal Omega. (B) Magnified view of J01-type deletion region of the alignment. Descriptions of the consensus symbols indicating the conservation level of amino acid residues are available at the Clustal Omega FAQ (https://www.ebi.ac.uk/seqdb/confluence/display/THD/Help+-+Clustal+Omega+FAQ). In brief, ‘:’ indicates residues that harbor a strongly similar physicochemical property to that in LQGH1, and ‘.’ indicates residues that harbor a weakly similar physicochemical property to that in LQGH1.

**S8 Fig.**
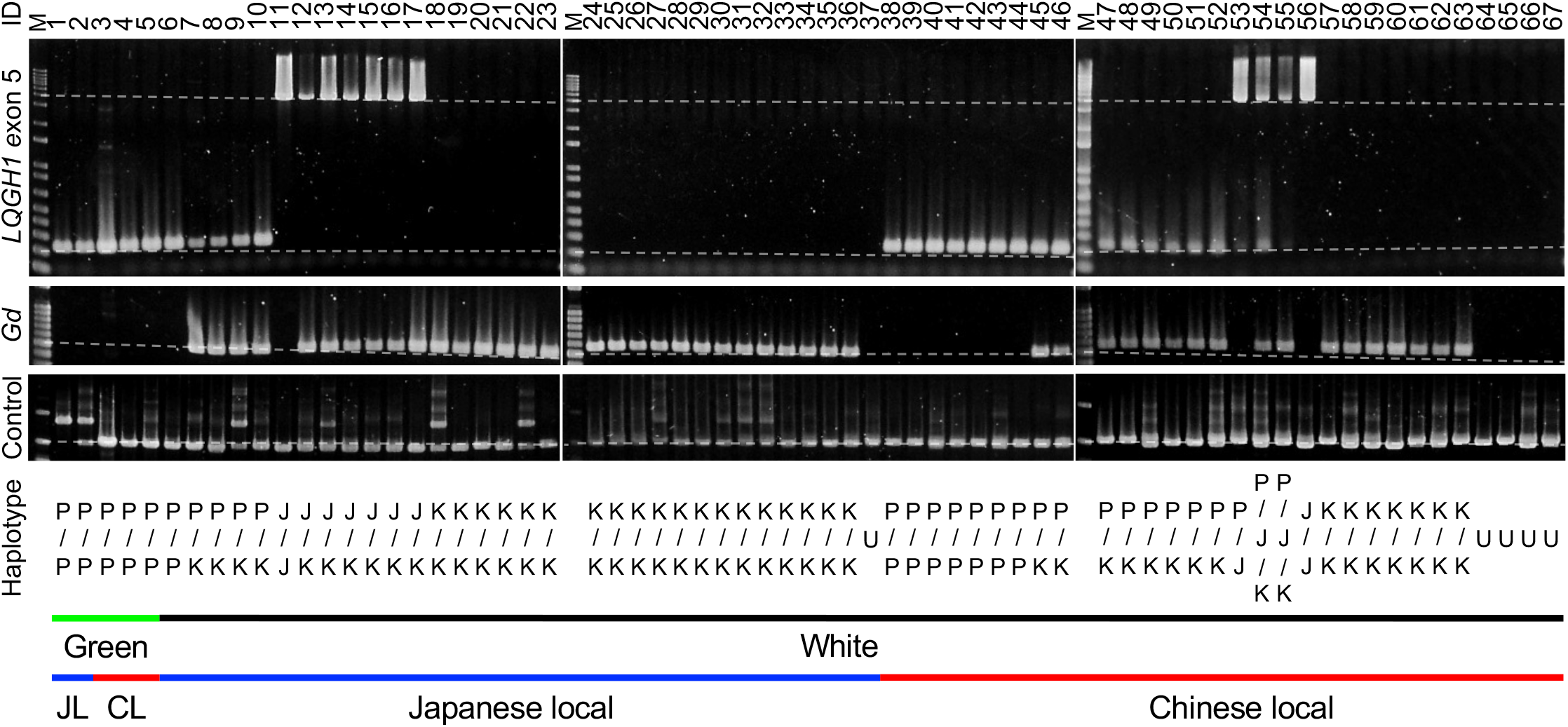
Genotyping of the Gd locus of Japanese or Chinese local white-cocoon strains. Haplotypes of the strains were determined by PCR genotyping. P/P, only the p50T genotype was detected; P/J/K, all p50T, J01, and Kosetsu genotypes were detected; J/J, only the J01 genotype was detected; J/K, both the J01 and Kosetsu genotypes were detected; K/K, only the Kosetsu genotype was detected; U, unknown, nothing detected. The primer set amplifying the genomic region from exons 1 to 2 of *KWMTBOMO14639* (*rp49*) was used as the control (predicted fragment length = 1548 bp). The grey dotted lines are straight lines connecting band markers of known size (M) that were applied at both ends of the samples. These lines represent sizes of 4000, 200, 500, and 1500 bp, from top to bottom. “JL” and “CL” stand for “Japanese local” and “Chinese local”, respectively. The colors indicate the color of the cocoon.

**S9 Fig.**
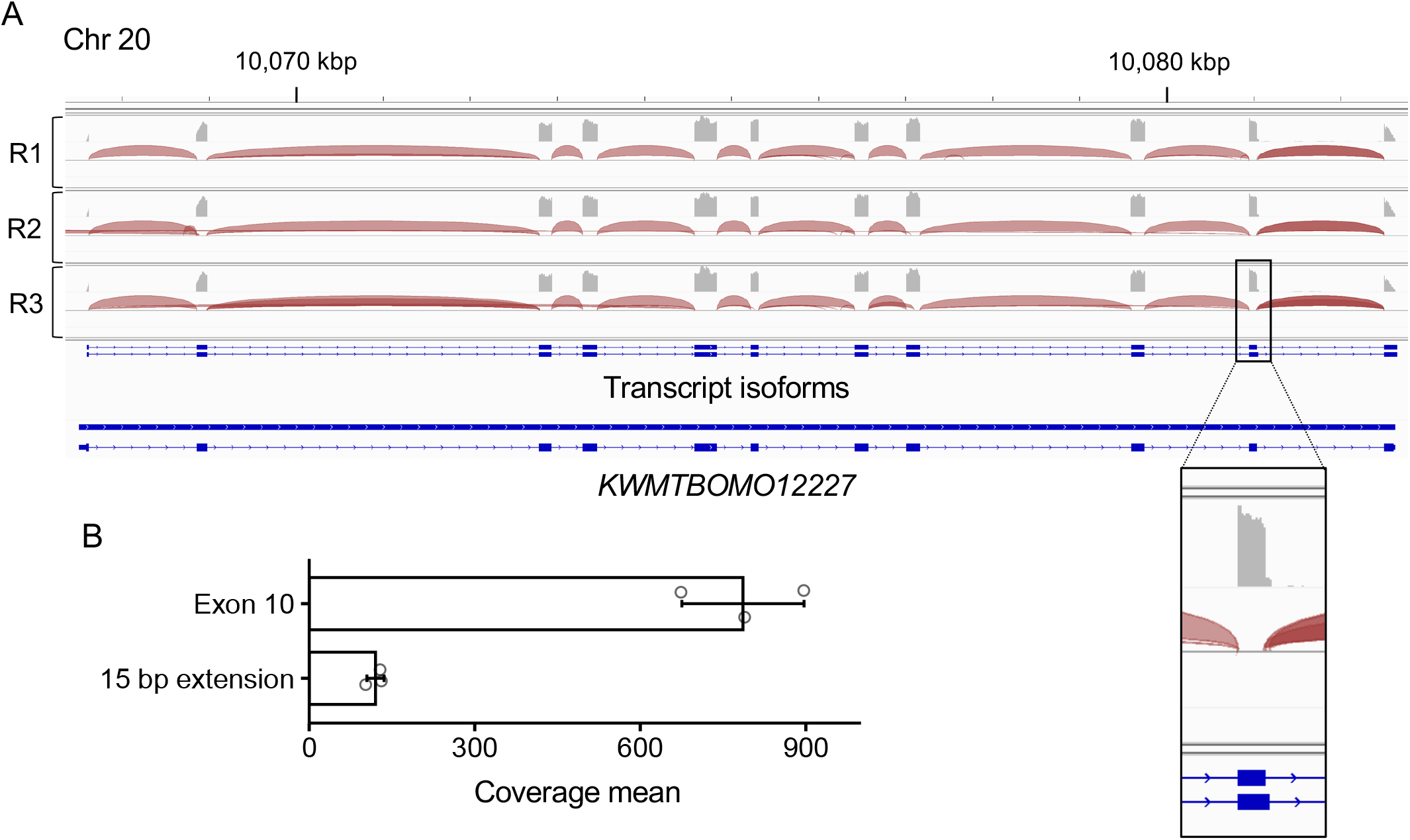
RNA-seq read alignment to the genomic sequence of *KWMTBOMO12227*. (A) Mapping result of the midgut-derived RNA-seq reads and *KWMTBOMO12227* transcript isoforms visualized with Integrative Genomics Viewer. Coverages on each exon and splice junctions of three replicates are illustrated on tracks R1–3. (B) Coverage mean of each base of exon 10 and the 15-bp extension region. Data are means ± SD.

**S10 Fig.**
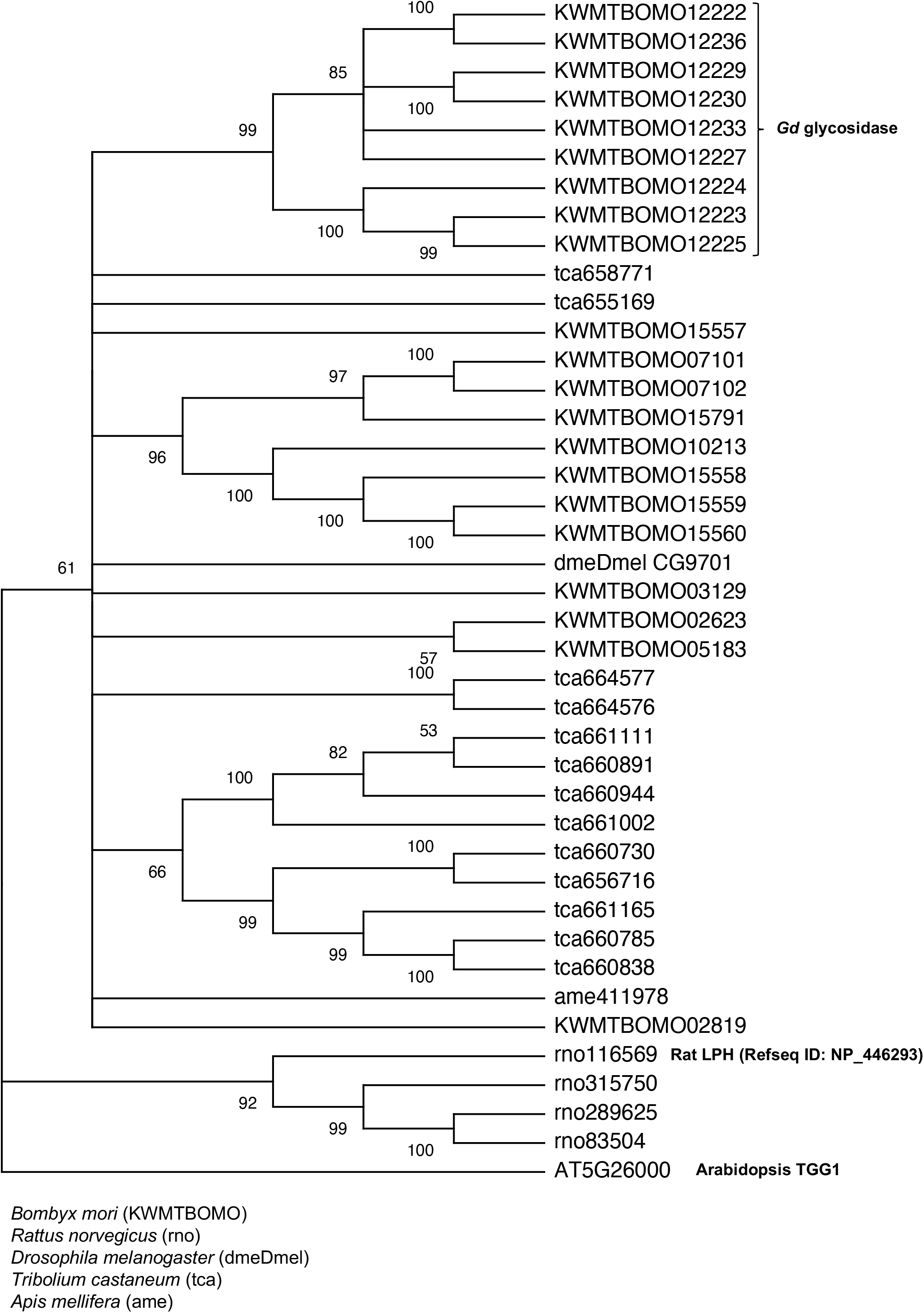
Phylogenetic tree of insect and rat proteins related to KWMTBOMO12227. Values on the nodes represent the bootstrap score (trials = 100). Unreliable nodes with bootstrap values under 50 are shown as multi-branching nodes. Arabidopsis thioglucoside glucohydrolase 1 (TGG1) was used as the outgroup.

**S11 Fig.**
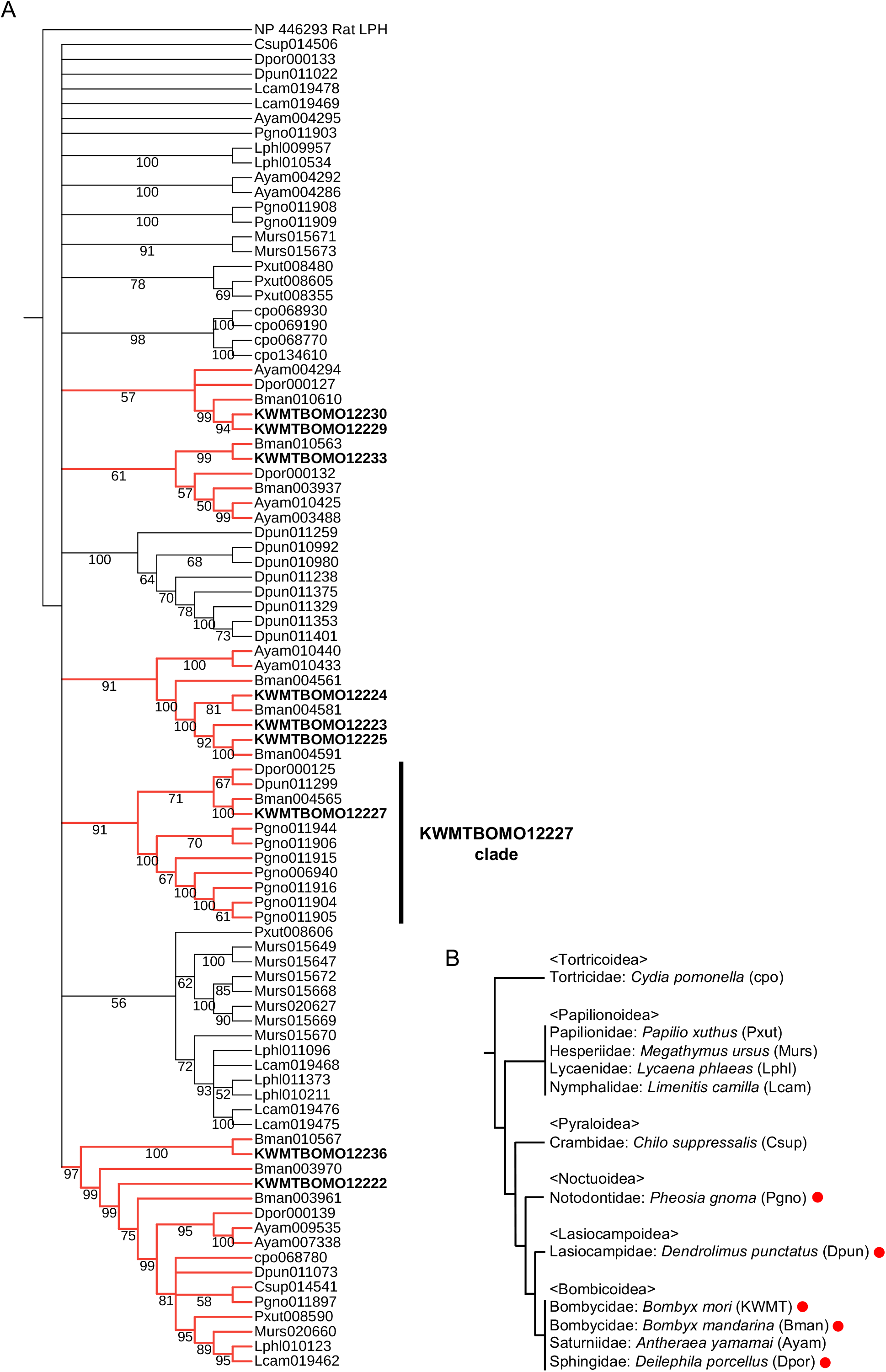
Phylogenetic tree of *Gd* glycosidase orthologs in Lepidoptera. (A) The phylogenetic tree was constructed by the maximum likelihood method. Values on the nodes represent the bootstrap score (trials = 100). Unreliable nodes with bootstrap values under 50 are shown as multi-branching nodes. Rat LPH was used as the outgroup. Red branches represent clades including the silkworm *Gd* glycosidases. Genes of the domestic silkworm are marked in bold. (B) Species tree of Lepidoptera with reference to the report by Kawahara et al.[71]. Red circles indicates species harboring an LQGH1 ortholog.

S1 Table. Information of the F2 individuals, parents, and phenotypic score used for the QTL analysis.

S2 Table. QTLs associated with flavonoid content in cocoon.

S3 Table. Composition of the semi-synthetic diets supplemented with quercetin or rutin. *Amount added per 100 g dry diet: K2HPO4, 2.25 g; CaCO3, 1.0g; KCl, 0.50 g; MgSO4, 0.3 g; FePO4・4H2O, 0.10g; ZnCl2, 0.01 g.

** This mixture includes cellulose powder (61.76 g/100 g). Amount added per 100 g dry diet: biotin, 0.2 mg; choline-HCl, 150 mg; folic acid, 0.2 mg; inositol, 200 mg; nicotinic acid, 10 mg, Ca-pantothenate, 15 mg; pyridoxine-HCl, 3 mg; riboflavin, 2 mg; thiamine-HCl, 2 mg.

*** Amount added per 100 g dry diet: propionic acid, 0.75 mL; chloramphenicol, 15 mg.

S4 Table. Summary of the Japanese and Chinese local silkworm strains examined.

S5 Table. Orthogroups of the 12 species of Lepidoptera and numbers of detected gene duplication events.

S6 Table. Domestic silkworm glycosidases.

S7 Table. Primers used in the experiments.

S1 Text. Determination and phylogenetic characterization of the amino acid sequence of KWMTBOMO12227.

