## Supplementary text for "A major endogenous glycosidase mediating quercetin uptake in *Bombyx mori*"

**S1 Text. Determination and phylogenetic characterization of the amino acid sequence of KWMTBOMO12227.**

### Transcript isoform and sequence of *KWMTBOMO12227*

Previously, our research group reported RNA-seq data for final instar larva tissues of the p50T strain [43]. By mapping the reads of the data derived from the midgut to the genomic sequence of the p50T strain reported by Kawamoto et al. [23], we determined the transcript isoform and the sequence of the gene modeled as *KWMTBOMO12227*. The sequence of reads aligned to *KWMTBOMO12227* was shown to be completely identical to the genomic sequence (S9A Fig). The genetic model accurately represented the dominant isoforms, but another isoform extending 15-bp downstream of the 10th exon was partially synthesized (S9A Fig). The termination codon in the extended region shortened the predicted protein sequence length of the minor isoform to 456 amino acids. The mean coverage of each base of the 15-bp extended region was only about 15% of that of the original 10th exon of *KWMTBOMO12227* (S9B Fig).

### KWMTBOMO12227 is a Macroheterocera-specific protein

To phylogenetically characterize the gene, we next collected homologous proteins from the domestic silkworm, the fruit fly (*Drosophila melanogaster*), the honeybee (*Apis mellifera*), the red flour beetle (*Tribolium castaneum*), and the rat (*Rattus norvegicus*) by BLASTp using the KWMTBOMO12227 protein sequence as a query to characterize the gene phylogenetically. This identified rat lactase/phlorizin hydrolase as a homologous protein. A phylogenetic tree of the proteins grouped the rat and insect proteins into different two clades and suggested that duplication of the *Gd* glycosidases has occurred after insect order divergence (S10 Fig). To investigate how KWMTBOMO12227 has evolved in Lepidoptera, a phylogenetic tree was constructed for lepidopteran-wide orthologous proteins of the domestic silkworm *Gd* glycosidases (S11A Fig). Ninety proteins from 12 lepidopteran insect species (Tortricidae, *Cydia pomonella*; Papilionidae, *Papilio xuthus*; Hesperiidae, *Megathymus ursu*s; Lycaenidae, *Lycaena phlaeas*; Nymphalidae, *Limenitis camilla*; Crambidae, *Chilo suppressalis*; Notodontidae, *Pheosia gnoma*; Lasiocampidae, *Dendrolimus punctatus*; Bombycidae, *Bombyx mori*; Bombycidae, *Bombyx mandarina*; Saturniidae, *Antheraea yamamai*; Sphingidae, *Deilephila porcellus*), classified in the same orthologous group as one of the nine silkworm *Gd* glycosidases using Orthofinder [37], were used in the analysis. The tree classified the *Gd* glycosidases into five clades: one clade including KWMTBOMO12227, one clade including KWMTBOMO12222 and KWMTBOMO12236, one clade including KWMTBOMO12223, KWMTBOMO12224 and KWMTBOMO12225, one clade including KWMTBOMO12229 and KWMTBOMO12230, and one clade including KWMTBOMO12233. The KWMTBOMO12227 clade included proteins found only in Macroheterocera (including Noctuoidea, Lasiocampoidea, and Bombycoidea). Of the investigated moth species in Macroheterocera, only the proteins of *Antheraea yamamai* were not included in the clade. Given the phylogenetic relationships of the lepidopteran superfamilies [71], this indicated that KWMTBOMO12227 has evolved through gene duplication after the divergence of Pyraloidea and Macroheterocera (S11B Fig). In the clade including KWMTBOMO12222 and KWMTBOMO12236, one or two orthologs of each of the 12 lepidopteran species were included, indicating that the gene evolved before the divergence of Tortricoidea and that the orthologs have an essential function for viability of lepidopteran insects.
